# A stable genomic variant for photoperiodic flowering plasticity to enhance grain mold escape and yield stability in sorghum

**DOI:** 10.64898/2026.04.01.715939

**Authors:** Djidjoho A.T. Hodehou, Cyril Diatta, Souleymane Bodian, Mbery Ndour, Diariétou Sambakhé, Bassirou Sine, Terry Felderhoff, Diaga Diouf, Geoffrey P. Morris, Ndjido A. Kane, Jacques M. Faye

## Abstract

Grain mold severely constrains sorghum [*Sorghum bicolor* (L.) Moench] productivity and grain quality in subhumid environments. Photoperiod-sensitive flowering plays a key role in mold avoidance and yield stability along north-south rainfall gradients. In response to the high susceptibility of elite cultivars in subhumid zones of Senegal, we developed and characterized a recombinant inbred line (RIL) population derived from Nganda (grain mold-susceptible) and Grinkan (photoperiod-sensitive) varieties. The population was evaluated across three distinct agro-ecological zones over two years. Environmental indices derived from genotype-environmental interactions, together with defined growth windows, strongly influenced flag leaf appearance (FLA), a photoperiodic flowering trait. Plasticity parameters (intercept and slope) for environmental indices, FLA, grain mold severity, and yield enabled identification of loci contributing to flowering response, mold resistance, and yield stability. The maturity gene *Ma1* and two QTLs for FLA, *qFLA6.2* and *qFLA6.3*, were identified, stable across environments, and colocalized with grain mold and yield QTLs. The wild-type *Ma1* allele from Grinkan delayed FLA and reduced grain mold damage but was not associated with increased yield. The *Ma1* effect was confirmed using the developed breeder-friendly KASP marker, Sbv3.1_06_40312464K, in 174 F_3_ three-way cross families. Photoperiod-sensitive lines with intermediate-to-late FLA alleles showed strong negative associations with mold damage. Overall, the identified stable loci and candidate lines provide foundations for effective molecular breeding of climate-resilient varieties.

**PLAIN LANGUAGE SUMMARY:** Grain mold is a fungal disease that reduces sorghum grain yield and quality, particularly in subhumid climates. With the limited number of resistant elite varieties, photoperiod-sensitive flowering to day length variation can contribute to grain mold escape at the end of rainy seasons. We characterized 286 sorghum recombinant inbred lines across three contrasting environments over two years along rainfall gradients in Senegal. Using flag leaf appearance (FLA), which is a photoperiodic flowering trait, strong genotype-environment interactions for FLA and genotypic plasticity were revealed. We identified and validated the common genomic locus associated with FLA variation and its plasticity across environments, the canonical maturity gene *Ma1*, which was influenced by temperature variation across environments. The presence of *Ma1* in the background of photoperiod-sensitive lines enhances grain mold avoidance and yield stability along rainfall gradients in Senegal.

**CORE IDEAS:**

- We investigated photoperiodic flowering plasticity in sorghum as a contributor to grain mold resistance and yield stability along rainfall gradients.
- The Maturity locus *Ma1* (*qFLA6.1*) is the major contributor of photoperiodic flowering and its plasticity across semi-arid and subhumid environments.
- Hybrid genotypes carrying two stable loci *qFLA6.1* and *qFLA6.2* sustain high grain mold avoidance in diverse environments.
- Photoperiod-sensitive lines with medium to late flowering times are effective in avoiding grain mold, while maintaining yield stability in subhumid regions.

## 1. INTRODUCTION

Breeding climate-resilient crop varieties has become a global priority under erratic rainfall and emerging diseases driven by climate variation. Among cereal crops, sorghum is one of the five most important cereals worldwide (Hossain et al., 2022) and famously resilience to broad environmental stresses, including heat, drought, and climate-related diseases makes it crucial for food and feed production (Harlan & de Wet, 1972). Sub-Saharan Africa accounts for about 71% of the global sorghum cultivation, where the grain is mainly used for food and provides 68% of dietary energy, along with protein (10%) and various oligo-elements (Dalton & Hodjo, 2020; FAO, 2024). However, grain mold reduces sorghum grain yield by up to 50 % during the rainy season (Audilakshmi et al., 2011). Sorghum grain mold affects seed and grain quality, limiting yield and market value (Aruna et al., 2012), but its genetic control is poorly understood. Improving sorghum grain mold resistance through developing climate-resilient genotypes is a crucial goal for sustainable breeding.

Grain mold refers to pre-physiological grain degradation that results from interactions of multiple opportunistic pathogenic fungal species (Ambekar et al., 2011), some of which produce potent mycotoxins that are harmful to human and animal health (Audilakshmi et al., 2011; Placinta et al., 1999) during grain development. Previous studies have shown that anthesis is the critical stage at which sorghum inflorescence is most susceptible to infection and colonization by fungi (Das et al., 2012; Little, 2000). The disease spreads further during caryopsis maturity until harvest especially under wet conditions (Audilakshmi et al., 1999, 2011). The use of exotic resistance sources to combat grain mold has faced challenges due to the disconnection between resistance traits and consumer preferences. For instance, a trait such as red pericarp, which may offer resistance, is less desirable by consumers (Audilakshmi et al., 2005; Bandyopadhyay et al., 2000; Diatta et al., 2019).

Advances in molecular genetics have facilitated the identification of genomic regions governing grain mold resistance in sorghum. Five Quantitative Trait Loci (QTL) linked to host incidence were mapped using a RIL population derived from RTx430/Sureño (Klein et al. 2001). Two other loci associated with grain mold resistance were identified using a sorghum mini-core collection (Upadhyaya et al., 2013). Recently, Cuevas et al. (2019) revealed three other loci associated with grain mold resistance in the US sorghum association panel (SAP). Although these findings provide potential tools and knowledge for breeding grain mold resistance, the genetic complexity of the trait across environments and genetic backgrounds hinder the practical use of these loci in breeding programs across regions (Aruna et al., 2021).

Photoperiod-sensitive varieties have been suggested as an effective strategy to avoid grain mold across environments (Bandyopadhyay et al., 2000). The ability of plants to modulate the onset of panicle initiation with varying sowing dates and photoperiod (day length) conditions is termed as photoperiodic flowering response (Bhosale et al., 2012). Photoperiodism allows sorghum to escape grain mold, as grain filling aligns with the onset of the dry season for a given zone, and minimizes the risk of mold development (Das et al., 2020), thus yield reduction. While flowering time itself is easy to measure, inferring photoperiod sensitivity across environments is difficult and time-consuming for breeders, especially at early generations (Guitton et al., 2018). Consequently, molecular markers are essential tools for tracking target traits such as photoperiod sensitivity in breeding (Zhao et al., 2018). In sorghum, photoperiod-sensitive flowering involves six major genes maturity genes, *Ma1 to Ma6* (Casto et al., 2019; Murphy et al., 2014). The dominant alleles confer delaying flowering under long days condition, with *Ma1* and *Ma6* both located on chromosome 6, have been reported to be the most impactful photoperiod-sensitive genes (Grant et al., 2023; Murphy et al., 2014). With the availability of cost-effective genotyping platforms such as KASP marker genotyping, nascent breeding programs can routinely monitor the alleles in segregating populations (Kumar et al., 2025; Maina et al., 2025; Rahman et al., 2023). However, to develop genotypes with broader and stable adaptation, it is crucial to understand the genetics and environmental contributions, and relationships between photoperiodic flowering, grain mold, and grain yield.

In this study, we used flag leaf appearance, which is a photoperiodic flowering-related trait and shown to be more precise than flowering time (Dingkuhn et al., 2008), to investigate the genetic control of photoperiodic flowering plasticity in sorghum in relation with grain mold and yield stability. We hypothesized the existence of common genetic control of photoperiodic flowering and its plasticity, where photoperiod sensitive alleles confer grain mold avoidance and grain yield stability. Moreover, at least one of the canonical flowering time loci would contribute to the north-south rainfall gradient adaptation in Senegal. To test this hypothesis, we investigated the plasticity of FLA using a RIL population evaluated in three contrasting locations over two years. The diurnal temperature range (DTR) from 49 to 59 days after planting was identified as the environment index that shapes the plasticity of FLA. QTL mapping studies using phenotypic mean values, alongside plasticity measures (i.e., intercept and slope) from environmental indexes, revealed common genetic control of the variation of target traits. (*iv*) Overall, we identified two stable loci, *Ma1* (*qFLA6.1*) and *qFLA6.2* harbored by photoperiod-sensitive lines, which can be used for grain mold reduction and climate-resilience breeding across environments.

## 2. MATERIALS AND METHODS

### 2.1. Plant materials and field trials

A biparental population was developed at the National Center of Agricultural Research of Senegal (CNRA) using elite inbred lines, Nganda and Grinkan. Grinkan is a Malian breeding line with moderate photoperiodic sensitivity and semi-open panicle shape (Kane et al., 2022). Nganda is a dual-purpose, grain mold susceptible and photoperiod-insensitive variety with farmers preferred traits such as white and tannin-free grain, high-yielding for food and sweet stem for feed (Diatta et al., 2021). The F_1_ hybrid plants were crossed to produce the F_2_ progeny, which was subsequently self-pollinated and advanced to produce 720 F_6:7_ families using the single seed descent method. For this study, 286 F_7:9_ families were randomly selected and evaluated along their parents during the cropping seasons of 2022 and 2023 in three locations in Senegal: Bambey, Sinthiou Maleme, and Sedhiou (Figure 1a, Table S1) using an alpha lattice design with two replications. Planting was done based on a two-row plot of 2 m long at a spacing of 0.7 m between rows and 0.2 m between plants (or hills) within a row. In 2022, a one-row plot of 4 m long each was used with 0.8 m between rows and 0.4 m between plants within a row due to the limited number of available seeds of F_6:7_ RILs. Plants were thinned to keep only one plant per hill, for a density of about 31,257 plants ha^-1^ and 143,000 plants ha^-1^ in 2022 and 2023, respectively. Standard agronomic practices for sorghum production were applied as described in Diatta et al. (2021).

**Figure 1:**
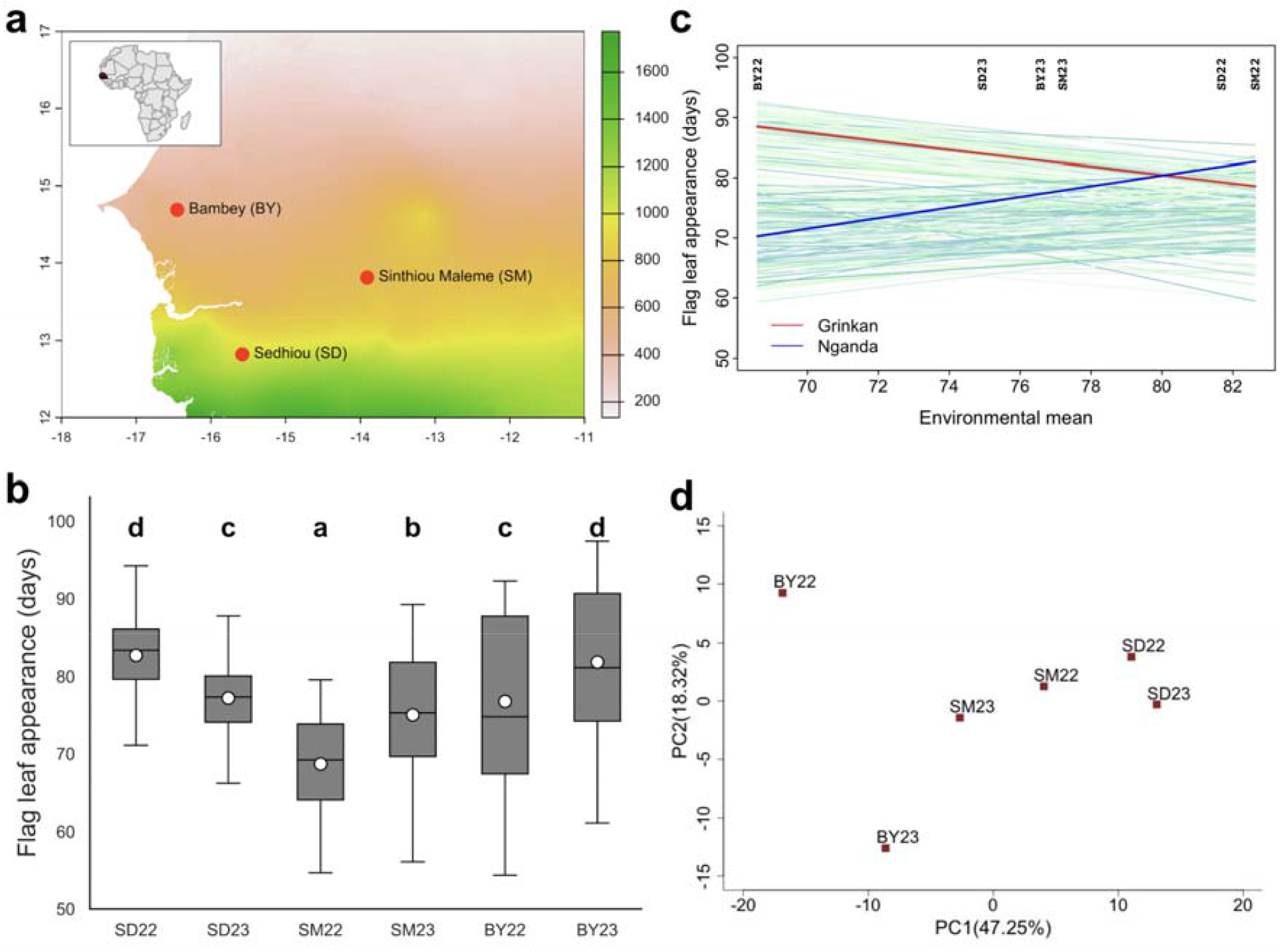
Genotype-environment interactions along rainfall gradient in Senegal. (a) The three contrasting locations along the precipitation gradient in Senegal used for phenotypic evaluation of the RIL population over two years, 2022 and 2023 under rainfed conditions, July to October. Contrasting locations are Bambey (BY), Sinthiou Maleme (SM) and Sedhiou (SD) over two years (2022 and 2023). The background color scale indicates the average annual precipitation in millimeters, with green representing the highest precipitation, pink representing the lowest precipitation, and yellow representing the intermediate zone. (b) Boxplot showing flag leaf appearance (FLA) variation in each environment. Mean values are represented as open circles within the boxplot. Multiple comparisons of FLA mean between environments are based on Tukey’s HSD test. FLA means sharing the same letter at the top of the boxplot are not significantly different. (c) Reaction norms of contrasting parents (Nganda, photoperiod insensitive and Grinkan, photoperiod sensitive) and their RIL progeny (n = 288) fitted with the joint-regression analysis using environmental means for FLA. (d) Principal component analysis of GxE effect for FLA based on the additive main effect and multiplicative interaction (AMMI) model across environments, BY22, BY23, SM22, SM23, SD22, and SD23.

A total of 174 F_3_ families derived from a three-way crossing were used to validate the effects of identified variants at *Ma1* in the current study. The first crosses were made between the five best grain mold resistance lines (Nganda x Sureño) and five best high-protein digestibility lines (Faourou x P721Q, an EMS mutant from Purdue University) of the Senegalese sorghum breeding program during the early hot off-season of 2022. The second cross was made between the produced F1s and six best photoperiod responsive lines during the early hot season of 2023 to establish a forward breeding scheme. Seeds from F_1_ plants were bulked and the F_2_ generation was grown through single-seed descent during the early hot off-season of 2024 at CNRA. The F_3_ panicles were threshed individually, and each was planted on a two-row plot of 2 m long at Sinthiou Maleme, following the same experimental design of 2023 season.

Six photoperiod-sensitive (n = 3) and insensitive (n = 3) RILs of the Senegalese sorghum breeding program, carrying the *Ma1* gene, were phenotyped for flag leaf appearance (FLA) along with two negative (Golobe and Nganda) and positive (Grinkan and Ni49) checks. The experiment was conducted at CERAAS from May to September 2024 in short day (SD; 8h light/16h dark) and natural long day (LD; day with > 13h daylight) conditions. This experiment was conducted to validate the effect of *Ma1*.

### 2.2. Phenotyping and data collection

FLA was recorded when half of the plants in a plot had their ligulated flag leaves visible, right before the booting stage. Grain yield (YLD; Kg. ha^-1^) were estimated on a plot basis, and scores for grain mold damage were recorded. The visual panicle (PGMR) and threshed (TGMR) grain mold ratings were recorded using the five points scale (1= highly resistant to 5= highly susceptible) described in Thakur et al. (2007). Following the geographic coordinates of each site, we retrieved daily rainfall precipitation (PRCP; mm day^-1^), maximum air temperature (TMAX; °C day^-1^), and minimum air temperature (TMIN; °C day^- 1^) from the database of the NASA POWER (Sparks, 2018) using the R package EnvRtype (Costa-Neto et al., 2021). In addition, the Daylength (DL) was calculated based on latitude using the *para_radiation* function of the R package EnvRType.

### 2.3. Statistical analyses

Statistical analyses were performed in R version 4.3.3 (R Core Team, 2025). Outliers were checked for all traits based on 1.5 × interquartile range (IQR) and removed using a R script custom by Chen et al. (2023). We used the “SpATS” package to correct phenotypic values for spatial trends (Rodríguez-Álvarez et al., 2018). The fitted values corrected for block and spatial effects were used in subsequent statistical analyses. The best linear unbiased estimator (BLUE) values were computed in the R package lme4 (Bates et al., 2015). The best linear unbiased predictor (BLUP) values for each trait were computed on a per-line basis from data of a given location across the two years or across all environments using the *ranef()* function within the lme4 R package by fitting the following model:

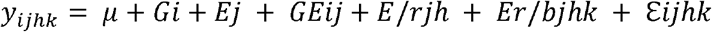

where *μ* is the observed value from each experimental unit, population mean, *Gi* effect of the i^th^ genotype, effect of the j^th^ environment, *GEij* interaction effect of the i^th^ genotype and the j^th^ environment, *E*/*rjh* effect of the h^th^ replicate nested to the j^th^ environment, *Er*/*bjhk* effect of the k^th^ block nested to the h^th^ replicate which is nested to the j^th^ environment, *ℰ ijhk* and the experimental error. Variance components were estimated based on the restricted maximum likelihood (REML) method (Patterson & Thompson, 1971). Broad-sense heritability (*H*^*2*^) of FLA in individual and combined environments was calculated as follows:

Individual environment:

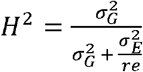

Combined environments:

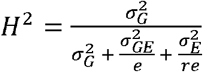

where *e*is the number of environments, *r* is the number of replicates per environment, and 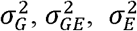 represents the variance of genotype, the interaction between the genotype and environment, and the residual error, respectively (Holland et al., 2002). Correlation analysis between traits was performed using BLUE values. The phenotypic plasticity of each line across environments was estimated using the Bayesian Finlay–Wilkinson regression model (FWR). In the FWR model, the observed phenotype of an individual in one environment can be expressed as:

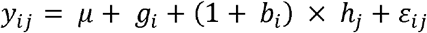

where *y*_*ij*_ is the phenotype of the *i*th line collected in the *j*th environment, *μ* is the regression intercept as population mean, *g*_*i*_ is the main effect of the *i*th line, *hj* is the environmental effect of the *j*th environment, 1+*b*_*i*_ represents the change in performance of the *i*th line per unit change in the environmental effect (*h*_*j*_) and *ε* _*ij*_ the residual error. All parameters, *g, b, h*, were treated as random effects, with a mean of zero and variance–covariance matrices □ following the previously published settings (Kusmec et al., 2017). Thus, *g*_*i*_ estimates the genotypic value as mean phenotype across environments. The value of 1+*b*_*i*_ measures the linear plasticity of a line *i* over environments, whereas (1+*b*_*i*_) × *h*_*j*_ indicates the phenotypic change of a line *i* in a given environment *j*. The Additive Main Effects and Multiplicative Interactions Model Addictive (AMMI) was used to understand the overall genotype-environment interaction. Genotype main effect (G), environment main effect (E) and GxE were analyzed by the AMMI model (Gauch & Zobel, 1990), represented by:

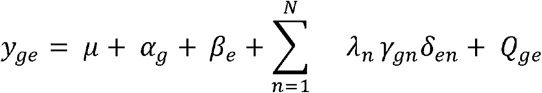

in which *y*_*ge*_ is the FLA mean of genotype g in environment *e, μ* is the grand mean, *αg* are the genotypic mean deviations (means minus grand mean), *β e* are the environmental mean deviations, *N* is the number of PCA axis retained in the adjusted model, *λ*_*n*_ is the singular value for PCA axis, *λ*_*n*_ *γ* _*gn*_ is the genotype eigenvector for PCA axis n, *δ* _*en*_ is the environment eigenvector for PCA axis *n,Q*_*ge*_ are the residuals, including AMMI noise and pooled experimental error. The expected distribution of *Q*_*ge*_ is normal. The PCA analysis level of significance was tested with the F-test according to Gollob (1968). The R scripts customized by Li et al. (2018) were adapted and used to implement the FWR and AMMI analyses.

We applied the CERIS algorithm (Li et al., 2018) to identify the combinations of environmental parameters and growth windows strongly correlated with the environmental mean. The mean value of each parameter was calculated for each window (different period) during development, followed by its correlation with the environmental mean of FLA. Each window started from a day (x) to a day (y) after planting. Six environmental indexes were evaluated: precipitation (PRCP), photoperiod expressed as day length (DL), temperature expressed as growing degree days (GDD), photothermal time (PTT = GDD x DL), photothermal ratio (PTR = GDD/DL), and Diurnal temperature range (DTR). GDD was defined as ((Tmax + Tmin)/2 – Tbase), where Tmax greater than 37.8°C was adjusted to 37.8°C, Tmin lower than 10°C adjusted to 10°C, which represents the sorghum base temperature (Tbase) (Borrell et al., 2020). The daily Diurnal temperature range (DTR) was performed as Tmax − Tmin, without any adjustment applied to Tmax or Tmin, such as GDD (Mu et al., 2022).

### 2.4 Genotyping and linkage map construction

Young leaves from the F_7_ RILs and parental lines were collected into 96-deep well plates according to the procedure required by the Intertek-Agritech laboratory. Samples were shipped to Diversity Arrays Technology in Australia through Intertek, Sweden, for DNA extraction and genotyping using DArTag technology containing a panel of 3,495 SNPs (Excellence in breeding, 2022). DArTag technology, which is a targeted genotyping-by-sequencing approach (Hardenbol et al., 2003) and data generation were performed following the methods described by Ongom et al. (2024). The genotypic data were filtered using VCFtools version 0.1.16 (Danecek et al., 2011) for low minor allele frequency (MAF > 1%), leaving 1,034 SNPs. All monomorphic markers (n = 189) and markers with more than 25% missing data were filtered out. Finally, 734 SNPs were used to construct a linkage map using the R package Onemap v 3.0.0 (Margarido et al., 2007). Pairwise recombination fractions were estimated considering a recombination fraction of < 0.5 and logarithm of the odds (LOD) score of 5.90, using default settings of the software.

### 2.5. QTL identification and candidate genes colocalization analysis

The QTL analysis was primary performed for FLA in each environment using the BLUE values. Intercept and slope values of FLA were obtained by regressing the phenotypes of each RIL on the environmental index. The intercept and slope values were then treated as phenotypes to identify QTL for the reaction-norm parameters (Figure 2d). QTL identification was conducted using the Composite interval mapping (CIM) method implemented with the software R/qtl (Broman et al., 2003). With α = 10%, LOD significance thresholds were determined based on 1000 permutations of the expectation-maximization algorithm. The QTL confidence interval spanned the genomic regions corresponding to 2-LOD drop from the peak. The “fitqtl” function in R/qtl was applied to determine the proportion of phenotypic variation explained (PVE) by each QTL. The QTL effect and interactions were generated using the “effectplot” function in R/qtl.

**Figure 2:**
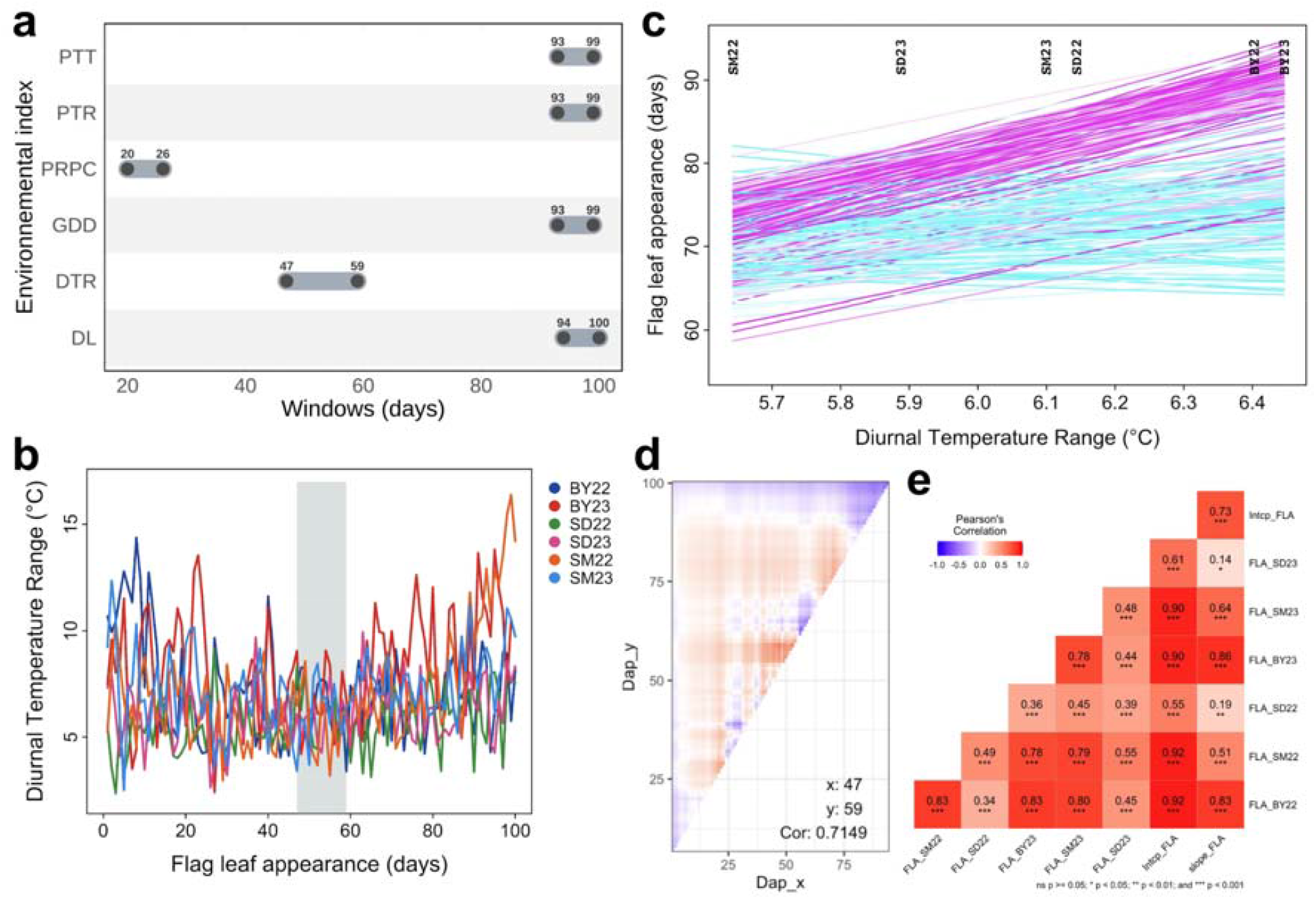
Determination of the critical window associated with environmental indexes for flag leaf appearance. (a) Critical window periods for recombinant inbred lines flag leaf appearance variation and environmental index combinations. (b) Diurnal temperature range (DTR) profiles across different environments. (c) Regression-fitted reaction norm using the environmental index (DTR_47–59_), between 47 and 59 days after planting (DAP) as the explanatory variable. Blue indicates lines with low slopes (low sensitivity), purple for lines with high slopes (high sensitivity), and white represents intermediate responses. (d) Heatmap of the correlation between the environmental index (any time window within the range 0-100 DAP, Dap) and the FLA phenotypic mean of the RIL population. (e) Pearson correlations between phenotypic means and plasticity parameters (intercept and slope) for FLA in different environments. Significance level is indicated as asterisks, (\**p* < 0.05; \*\**p* < 0.01; \*\*\**p* < 0.001).

QTL names were generated by starting with the letter “q” to indicate QTL, followed by the abbreviation of the trait name, the chromosome number, and an index number for each QTL (e.g., *qFLA6.1* denotes the first FLA QTL on chromosome 6). QTL for YLD, PGMR, and TGMR were also identified to assess colocalizations with FLA QTL. All identified QTL were further compared with previously known flowering genes to explore potential candidate genes.

### 2.6. QTL validation and origin of the photoperiod-insensitive allele at *Ma1*

To validate the effect of the QTL at *Ma1* gene, the developed Kompetitive allele-specific PCR (KASP) marker at the causal variant Sbv3.1_06_40312464K in *Ma1* was used to genotype the 174 F_3_ individuals phenotyped for FLA in natural field. In addition, the Sbv3.1_06_40312464K marker was used to select three photoperiod-sensitive and photoperiod-insensitive lines from the Senegalese breeding program, with two negative and positive photoperiod checks, which were evaluated under short-day and long-day conditions in greenhouse. Two foliar leaf discs were collected from 3-weeks old individual plants for Genomic DNA extraction and low-density KASP genotyping at Intertek Agritech Lab in Sweden. The *cross_qc_heatmap* function implemented in the panGenomeBreedr R package (Kena et al., 2025) was used to analyze the genetic background of parents and the selected RIL progenies. The Africa map showing the origin of the photoperiod-insensitive allele at *Ma1* was designed following a model from Klein et al. (2015) and using a custom script in ggplot2.

## 3. RESULTS

### 3.1. Phenotype variation and genotype-environment interactions

To determine whether the phenotypic data quality was good for genetic dissection of photoperiodic flowering plasticity, we assessed the genetic contribution of RILs to phenotypic variation of flag leaf appearance (FLA) across environments. FLA exhibited abundant phenotypic variation and low coefficient of variation (CV, values ranging from 6.73% in SD23 to 14.78% in BY22) across all environments (Figure 1b, Table 1). The RILs showed transgressive segregation in all environments, except in SM22, where the female parent, Nganda, had a larger FLA value (96 days) than the maximum FLA (86 days) of the RILs. The broad-sense heritability (*H*^2^) of FLA in each environment was generally high (Table 1), except in SD22 and SD23, which exhibited moderate (SD22; *H*^2^ = 0.59) and low (SD23; *H*^2^ = 0.24) heritability, respectively. The broad-sense heritability across the six environments was high (*H*^2^ = 0.89), based on BLUP values. Also, the correlation between FLA and days to flowering showed a strongly significant positive correlation (*r = 0.99; p < 0.0001*) across all environments.

**Table 1:** Summary statistics for flag leaf appearance in six environments and based on BLUP values.

| Trait† | Env‡ | Grinkan | Nganda | *RIL population statistics* |  |  |  |  |  |  |
| --- | --- | --- | --- | --- | --- | --- | --- | --- | --- | --- |
|  |  |  |  | Min | Max | Mean ± SD | §CV (%) | Skewness | Kurtosis | ¶H<sup>2</sup> |
| FLA | BY22 | 87 | 66 | 43 | 95 | 76 ± 11 | 14.78 | -0.06 | -1.39 | 0.87 |
|  | BY23 | 95 | 78 | 59 | 100 | 82 ± 10 | 12.12 | -0.21 | -1.07 | 0.84 |
|  | SM22 | 86 | 96 | 52 | 86 | 68 ± 6 | 9.37 | -0.22 | -0.72 | 0.89 |
|  | SM23 | 78 | 84 | 51 | 92 | 74 ± 9 | 11.56 | -0.34 | -0.77 | 0.87 |
|  | SD22 | 73 | 67 | 65 | 99 | 83 ± 7 | 8.68 | -0.10 | 0.21 | 0.59 |
|  | SD23 | 80 | 79 | 79 | 90 | 77 ± 5 | 6.73 | -0.10 | -0.42 | 0.24 |
|  | BLUP | 80 | 74 | 62 | 85 | 74 ± 6 | 7.53 | -0.08 | 1.86 | 0.89 |

† FLA, flag leaf appearance; ‡ Env, environment; § CV, coefficient of variation; ¶Broad- sense heritability; SD, standard deviation from the mean.

Subsequent variance component analysis indicated high genotypic variance (34%) for FLA, followed by environmental variance (28%), and the interactions between genotype and environment (14%) (Table S2). GxE was partitioned into the heterogeneity of slope (64%) and error (36%) (Table S3). The FLA environmental mean of the entire population varied from 69 DAP (days after planting) in BY22 to 83 DAPs in SM22 (Figure 1c). Following the additive main effects and multiplicative interaction (AMMI) models, the first two principal components accounted for 65.57% of GxE, and the plot based only on GxE effects showed BY22 as the environment with the highest GxE effects (Figure 1d, Table S4).

### 3.2. Critical windows and environmental indices

We determined the critical environmental factor that influenced the variation of the FLA using the CERIS model. The average values of the diurnal temperature range from 47 to 59 DAP (DTR_47-59_; *r = 0.72*) showed a stronger correlation with the environmental mean than the average values of other environmental factors within various growth windows (Figure 2a,b,c). Within the period spanning 100 DAP, precipitation (PRCP_20-26_; *r = 0.68*) was more correlated with the environmental mean at the early vegetative stage of plants, whereas photothermal time (PTT_93-99_; *r = 0.69*), photothermal ratio (PTR_93-99_; *r = 0.67*), growing degree days (GDD_93-99_; *r =0.68*), and day length (DL_94-100_; *r = 0.5*) were higher at the end of growth stage of plants (Figure 2a). In addition, between 30 and 80 DAPs, both day length and photothermal time showed a moderate correlation (*r = 0.44*) with the environmental mean. We defined the regression-adjusted reaction norms (intercept and slope), based on the environmental index DTR_47-59_ as explanatory variables, which were modeled for all progenies (Figure 2d). Pearson correlations between FLA measured in the six environments and the reaction norm parameters were all significantly positive (Figure 2e). The environment indices were used instead of the environmental mean to determine the genetic control of FLA in relation to environmental variation.

### 3.3. Genetic control of flag leaf appearance along precipitation gradient

To identify putative genomic variants that contribute to the genetic variation of FLA across the north–south precipitation gradient, we conducted a CIM using the phenotypic mean values of the RIL population. A total of 734 high confidence markers were used to construct the linkage map out of the 3,495 markers of the sorghum DArTag panel. Among these markers, 11 causal variants KASP-based markers at the canonical maturity loci, *Ma1* (n = 8, each specific to the 8 different alleles of the gene), *Ma2* (n = 1), *Ma3* (n = 1), and *Ma6* (n = 1) are included. A total of thirteen (13) QTLs (*qFLA1.1*; *qFLA1.2*; *qFLA1.3; qFLA2*; *qFLA3.1*; *qFLA3.2*; *qFLA5.1*; *qFLA6.1*; *qFLA6.2*; *qFLA6.3*; *qFLA7.1*; *qFLA10.1; qFLA10.2*) were detected using FLA values in each environment (Figure S1, Table 2, Table S5). The proportion of phenotypic variance explained (PVE) by these QTLs ranged from 0.5% to 56.2% (Table 2, Table S5). The QTL (*qFLA6.1*) located at 40.31 Mbp on chromosome 6, which corresponds to one of the causal variants at the known maturity gene *Ma1*, formerly known as *PSEUDO-RESPONSE REGULATOR7* (*SbPRR7*), was identified in two semi-arid environments, BY22 and BY23. The functional allele *SbPRR7-2* (photoperiod-sensitive) was identified in this genetic population and accounts for 56.2% and 45% of the PVE in BY22 and BY23, respectively.

**Table 2:** The top common QTLs identified for FLA, plasticity measures of FLA, PGMR, TGMR, and YLD in contrasting environments.

| Trait | Env<sup>a</sup> | Chr<sup>b</sup> | QTL | Pos (Mbp)<sup>c</sup> | Pos (cM)<sup>d</sup> | CI (cM)<sup>e</sup> | LOD<sup>f</sup> | PVE<sup>g</sup> | Effect | Allele<sup>h</sup> |
| --- | --- | --- | --- | --- | --- | --- | --- | --- | --- | --- |
| FLA | BY22 | 2 | *qFLA2* | 47.15 | 23.6 | 12–30 | 6.9 | 13 | -4 | G |
|  |  | 6 | *qFLA6.1* | 40.31 | 119.3 | 119–120 | 34.3 | 56 | -8 | G |
|  | BY23 | 6 | *qFLA6.1* | 40.31 | 119 | 110–120 | 21.3 | 45 | -6 | G |
|  | SD22 | 6 | *qFLA6.2* | 1.83 | 132 | 128–132 | 11.6 | 20 | -3 | G |
|  | SD23 | 6 | *qFLA6.2* | 1.83 | 132 | 128–132 | 11.4 | 19 | -2 | G |
|  | SM22 | 2 | *qFLA2* | 47.15 | 75.3 | 71–80 | 3.6 | 6 | 2 | N |
|  |  | 6 | *qFLA6.3* | 42.12 | 118.5 | 128–132 | 12.2 | 38 | -4 | G |
|  | SM23 | 3 | *qFLA3.2* | 1.23 | 556 | 328–557 | 4.8 | 10 | 3 | G |
|  |  | 6 | *qFLA6.3* | 42.12 | 118 | 110–120 | 14.5 | 29 | -5 | G |
|  | Intercept | 3 | *qFLA3.2* | 1.23 | 556 | 20–557 | 3.6 | 3 | 1 | G |
|  | Intercept | 6 | *qFLA6.3* | 42.12 | 118 | 110–132 | 16.9 | 45 | -4 | G |
|  | Slope | 6 | *qFLA6.1* | 40.31 | 119.3 | 119–120 | 46 | 46 | -6 | G |
| PGMR | BY22 | 4 | *qPGMR4* | 59.55 | 67.2 | 66–78 | 5.6 | 0.5 | -0.03 | N |
|  | SD22 | 6 | *qPGMR6.2* | 1.83 | 131.8 | 128–132 | 7.3 | 15 | 0.2 | G |
| TGMR | BY22 | 4 | *qTGMR4* | 59.55 | 99.2 | 66–78 | 3.7 | 6 | 0.2 | G |
|  | SD22 | 6 | *qTGMR6.1* | 1.83 | 132 | 128–132 | 7.9 | 12 | 0.3 | G |
|  | SM22 | 6 | *qTGMR6.2* | 42.12 | 118.5 | 110–126 | 12 | 18 | 0.4 | G |
| YLD | SM23 | 6 | *qYLD6.2* | 40.31 | 119 | 110–126 | 14 | 19 | 454 | N |

FLA: Flag leaf appearance; PGMR, panicle grain mold rate at physiological maturity; TGMR, grain mold rate after threshing; YLD, grain yield; <sup>a</sup>Environment; BY, Bambey site; SD, Sedhiou site; SM, Sinthiou Maleme site in 2022 and 2023; <sup>b</sup>Chromosome; <sup>c</sup>Physical position (Mbp); <sup>d</sup>Genetic map position in centiMorgans; <sup>e</sup>2.0-LOD confidence interval; <sup>f</sup>Log10 likelihood ratio comparing full model with reduced model; <sup>g</sup>Proportion of phenotypic variation explained; <sup>h</sup>Source of the allele with the favorable effect; G, Grinkan; N, Nganda. The favorable alleles were identified according to the breeding objective of each target trait. Higher values for FLA and YLD, and lower values for PGMR and TGMR, were considered favorable.

Another QTL (*qFLA6.2*) located at 1.83 Mbp on chromosome 6 was identified in the most humid environments, SD22 and SD23. This QTL accounted for 19% of the PVE. The allele of the parental line, Grinkan (G allele) at *qFLA6.1* and *qFLA6.2* was associated with late flag leaf appearance, relative to the photoperiod-insensitive allele of the parental line, Nganda (Table 2). The QTL, *qFLA6.3*, located at 42.12 Mbp on chromosome 6, was identified in both subhumid environments, SM22 and SM23. This QTL explained approximately 30 to 38% of PVE (Table 3). The peak on chromosome 2 (*qFLA2.1*) with significant effect (10-10.3%) was identified in two humid environments (SD22 and SM23). To characterize the genetic control of phenotypic plasticity, we used the computed environment reaction norms based on phenotypic values and environment variables. Five putative plastic QTLs were identified based on the intercept and slope values (Table 2, Table S5). Comparing these QTLs with those found based on phenotypic mean values in each environment, only one novel QTL (*qFLA10.2*; PVE, 0.8%, effect size, 0.6) was identified on chromosome 10 (Table S5).

**Table 3:** Pearson correlation between allelic classes of stable QTLs for FLA with grain mold damage and yield reduction.

|  | *Ma1(qFLA6.1)* |  |  | *qFLA6.2* |  |  |
| --- | --- | --- | --- | --- | --- | --- |
|  | *FLA time in Bambey* |  |  | *FLA time in Sedhiou* |  |  |
| *Trait*† | Early | Medium | Late | Early | Medium | Late |
| PGMR | **-0.27\*** | **-0.58\*\*\*** | **-0.48\*\*\*** | **-0.50\*\*\*** | **-0.52\*\*** | **-0.44\*\*\*** |
| TGMR | 0.05ns | -0.03ns | 0.04ns | **-0.44\*\*\*** | **-0.57\*\*** | **-0.42\*\*\*** |
| YLD | -0.19ns | -0.33ns | -0.15ns | 0.18ns | -0.007ns | **-0.22\*** |
|  | *FLA time in Sedhiou* |  |  | *FLA time in Bambey* |  |  |
| PGMR | **-0.34\*\*\*** | -0.22 | -0.12 | **-0.31\*\*\*** | -0.19ns | **-0.18\*** |
| TGMR | **-0.61\*\*\*** | **-0.57\*\*\*** | **-0.49\*\*\*** | **-0.48\*\*\*** | **0.57\*\*\*** | **-0.41\*\*\*** |
| YLD | 0.11ns | 0.33ns | 0.14ns | 0 | 0.25ns | 0.12ns |

† PGMR, Panicle grain mold rating; TGMR, threshed grain mold rating; YLD, grain yield based on BLUE values from Bambey and Sedhiou in 2022 and 2023 for qFLA6.1 and qFLA6.2, respectively; \*p < 0.05; \*\*p < 0.01; \*\*\*p < 0.001; ns: not significant.

### 3.4. Colocalization and epistatic interactions of QTL and known genes

We mapped QTLs for grain mold and yield to assess colocalizations with FLA QTLs. Out of these loci, *qFLA6.2* and *qFLA6.3* colocalized with PGMR and TGMR QTLs, whereas *qFLA6.1* colocalized with the YLD QTLs. *qPGMR6.2* and *qTGMR6.1* explained a large proportion of the PVE (14.8% and 11.8%, respectively) and colocalized with *qFLA6.2* in the same environment (SD22). *qTGMR6.2* accounted for the largest PVE (17.8%) and colocalized with *qFLA6.3* (Table 2). Across these QTLs, the Grinkan parental allele consistently reduced mold damage and was therefore considered the favorable allele (Figure S2, Figure S3). In the SM23 analysis, the largest YLD QTL colocalized with the largest FLA QTL at *Ma1*. However, the Grinkan allele was not associated with increase grain yield (Figure S4). Colocalization analysis revealed that FLA QTL overlapped with five known photoperiod-related flowering genes, namely *GI, CN12, Ma1, LHY*, and *CO* (Figure 3a). Grain mold (GMD) QTLs colocalized with four photoperiod-related florigen genes (*Ehd1, CN12, Hd1*, and *Ma1*), whereas YLD QTL colocalized exclusively with *Ma1*.

**Figure 3:**
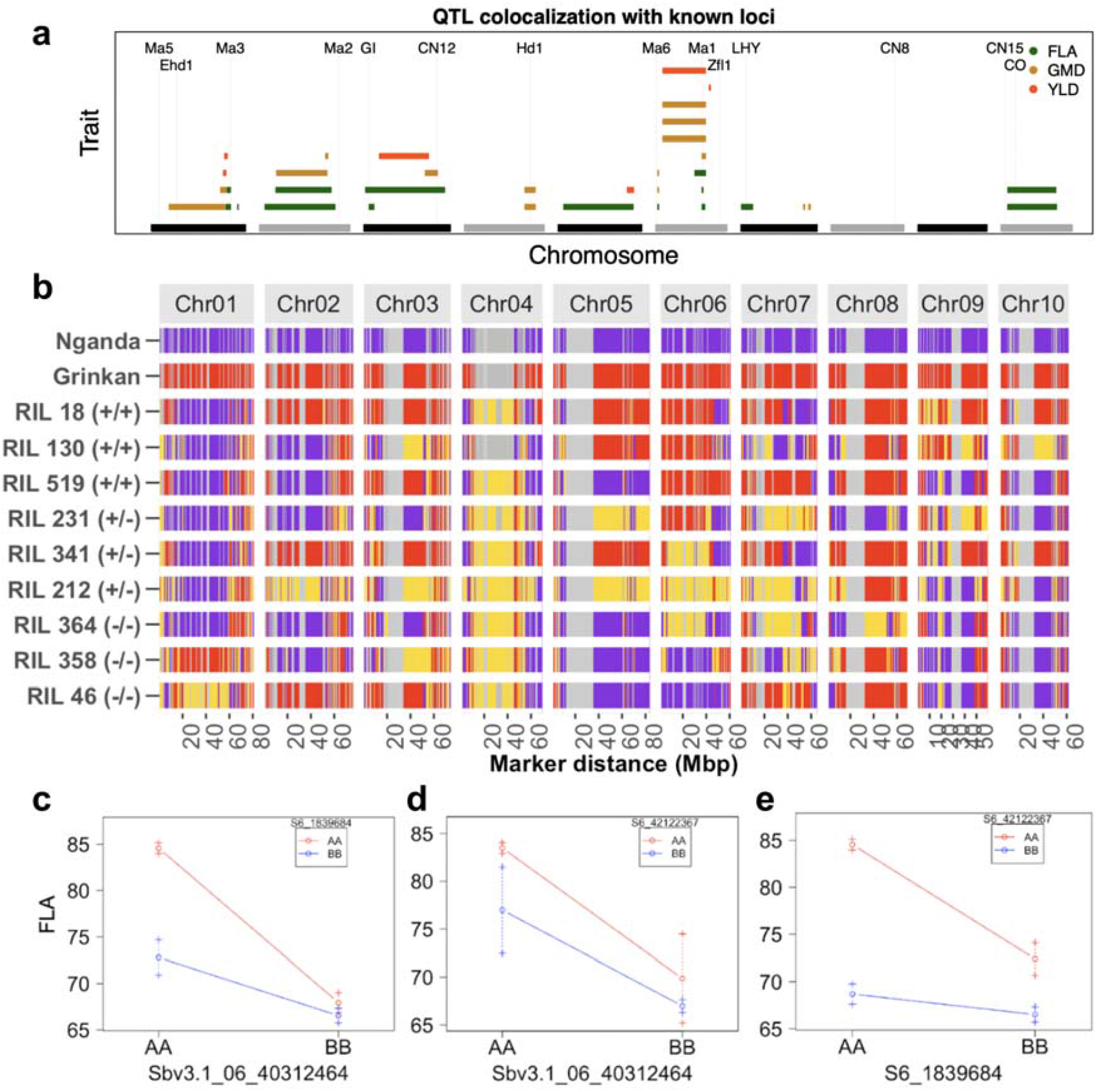
Common quantitative trait loci (QTLs) associated with flag leaf appearance, grain mold and yield. (a) Colocalization of the top associated QTLs for flag leaf appearance FLA, grain mold (GMD) and grain yield (YLD) with known candidate photoperiodic flowering time genes. Genomic positions of known candidate genes are indicated with vertical dashed lines. (b) Heatmap comparing the genetic background of the parents and three classes of selected recombinant inbred line (RILs), based on agronomic performance and allelic class of the QTL at *Ma1*, with genotypes with the homozygous dominant allele indicated as (+/+), homozygous recessive allele as (−/−), and heterozygous genotypes as (+/−), across all markers. The purple color indicates Nganda background, the red color indicates Grinkan background, the yellow color indicates heterozygous genotypes, and gray indicates monomorphic sites. (c-e) Two-way interaction plots for genotypic effects of the top three QTLs for FLA, (c) *Ma1* (*qFLA6.1*) vs. *qFLA6.2*, (d) *Ma1 vs. qFLA6.3*, and (e) *qFLA6.2 vs. qFLA6.3*.

We hypothetically classified QTLs alleles as photoperiod sensitive (PPS) or photoperiod insensitive (PPI) into late or early genotype class, respectively, including their intermediate heterozygote class, relative to FLA; and further examined the genome-wide background of selected RILs representing each allelic class with PPS (annotated +/+), PPI (−/−), and intermediate heterozygous (+/−). All individuals showed a mosaic genome inherited from both parents (Figure 3b). To determine the epistatic interaction effects among alleles at photoperiodic flowering-associated loci, we focused on the top three significantly associated QTLs, with putative major effect size, *qFLA6.1* (*Ma1*), *qFLA6.2*, and *qFLA6.3* on chromosome 6 (Table S6). A significant interaction between *Ma1* and *qFLA6.2* was observed, with a LOD score of 5.3, explaining 3.3% of the phenotypic variation of the FLA (Figure 3c, Table S6). *Ma1* × *qFLA6.3* interaction was significant with a LOD score of 4.8 and explained 3% of phenotypic variation (Figure 3d, Table S6). Significantly low interaction was detected between *qFLA6.2* × *qFLA6.3* (Figure 3e, Table S6).

### 3.5 Environmental stability of major QTLs across agroecological zones

At the positions of *Ma1, qFLA6.2*, and *qFLA6.3* on chromosome 6, the genotypes aligned with their corresponding genetic background and allelic class, homozygous to the Grinkan allele for the PPS class, homozygous to the Nganda allele for the PPI class, and heterozygous for the intermediate class (Figure 4a). We identified 152 RILs classified as late maturity lines that harbor the functional allele at *Ma1*, and 103 early maturity RILs with the recessive allele at *Ma1* (Figure 4b), which explained 46% of PVE across the environments (slope).

**Figure 4:**
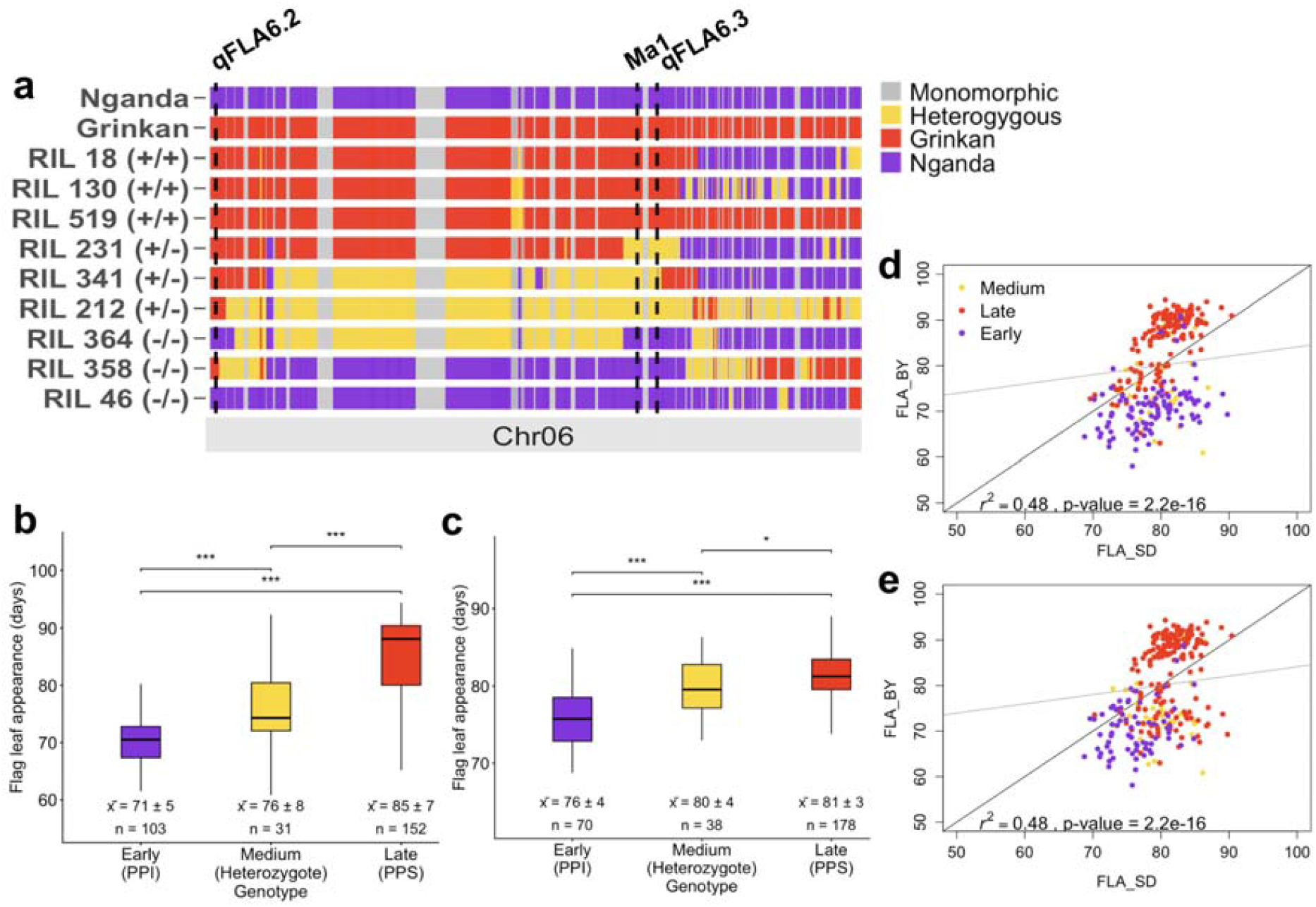
Allelic associations of top three QTLs with photoperiodic flowering variation. (a) Local heatmap with the annotation of the position of *Ma1, qFLA6.2*, and *qFLA6.3* on Chromosome 6. The different colors represent the genetic background of the parents (purple for Nganda, red for Grinkan) and three classes of selected recombinant inbred line (RILs) based on agronomic performance and allelic state at *Ma1* locus, for homozygous dominant (+/+), homozygous recessive (−/−), and heterozygous (+/−) progenies across all markers. (b,c) Associations of the allelic classes of the top two associated QTLs, (b) *qFLA6.1* or *Ma1* and (c) *qFLA6.2*, with FLA variation, based on BLUE values and two-tailed *t*-tests. The asterisks *** denotes *p* < 0.0001; **, *p* < 0.001; *, *p* < 0.05; and *ns*, not significant. (d,e) One-on-one ratio correlation plots for FLA in Sedhiou (SD) and Bambey (BY) most contrasting environments for *Ma1* and *qFLA6.2*, respectively. Each point represents a mean phenotypic value of a RIL, color-coded by allelic classes as early, medium, late FLA for each QTL.

Significant differences were observed between genotypic classes for the top two loci (Figure 4b,c). Next, we next assessed the consistency of the effects of the top two loci, *Ma1* and *qFLA6.2*, across two contrasting environments (SD and BY), based on BLUE values. Correlation analyses revealed that both loci exhibited stable effects on FLA under the environments. A moderate but highly significant correlation was observed between FLA values in SD and BY for both *Ma1* and *qFLA6.2* (0.48, *p < 2.2e-16*), with most individuals distinctly grouped by flowering class (Early, Medium, Late) (Figure 4d,e).

### 3.6. Effects of time of flag leaf appearance on grain mold damage and yield variation

To characterize the relationship between the homozygous allelic classes at the top two loci, with grain mold damage and grain yield reduction, we use PGMR, TGMR, and YLD based on BLUE values from BY and SD for *Ma1* and *qFLA6.2*, respectively. Significant associations were observed between allelic classes at the two QTLs with grain mold damage and grain yield (Table 3). All allelic classes at QTLs had significantly negative correlation with PGMR, with medium and late classes showing the strongest associations at *Ma1* (−0.27, *p <* .05; −0.58, *p <* .001; −0.48, *p <* .001 for early, medium and late classes, respectively), while early and late classes showing the strongest associations at *qFLA6.2* (−0.5, *p <* .001; −0.52, *p <* .01; −0.44, *p <* .001, respectively). There was no significant association of allelic classes at *Ma1* with TGMR (at *p <* .05). Whereas significant negative associations were observed between all allelic classes at *qFLA6.2* and TGMR (−0.44, *p <* .001; −0.57, *p <* .01; −0.42, *p <* .001 for early, medium, and late classes, respectively), with the medium class having the least significant association. The late allelic class only at *qFLA6.2* was mildly associated with YLD (−0.22, *p* < .05).

### 3.7. Stable QTL alleles confirmation

The trait predictiveness of the KASP-based causal marker, Sbv3.1_06_40312464K, which was developed for one of the allele of the gene *Ma1*, was accessed using the pot experiment under managed short- and long-day conditions in greenhouse (Figure 5a) and field experiment. All three selected lines and parent Grinkan, with PPS allele G had significantly lower FLA means under SD condition than under LD condition in greenhouse (Figure 5b). There was no effect of day length variation on the lines with PPI allele T and parent Nganda. The 174 individual F_3_ families, phenotyped under long days rainfed field conditions, were divided into three groups based on the KASP marker alleles (Figure 5c). Group 1 carried the dominant PPS allele, *Ma1* (GG genotype), group 2 carried the PPI allele *ma1* (TT genotype), and group 3 was heterozygote (Figure 5d). Individuals with the G allele had a significantly (*p <* .0001) higher mean FLA (70 DAPs) than those with the T allele (65 DAP) (Figure 5d). Likewise, individuals with the G allele had significantly (*p* < .01) higher FLA mean value than the heterozygous individuals (66 DAP).

**Figure 5:**
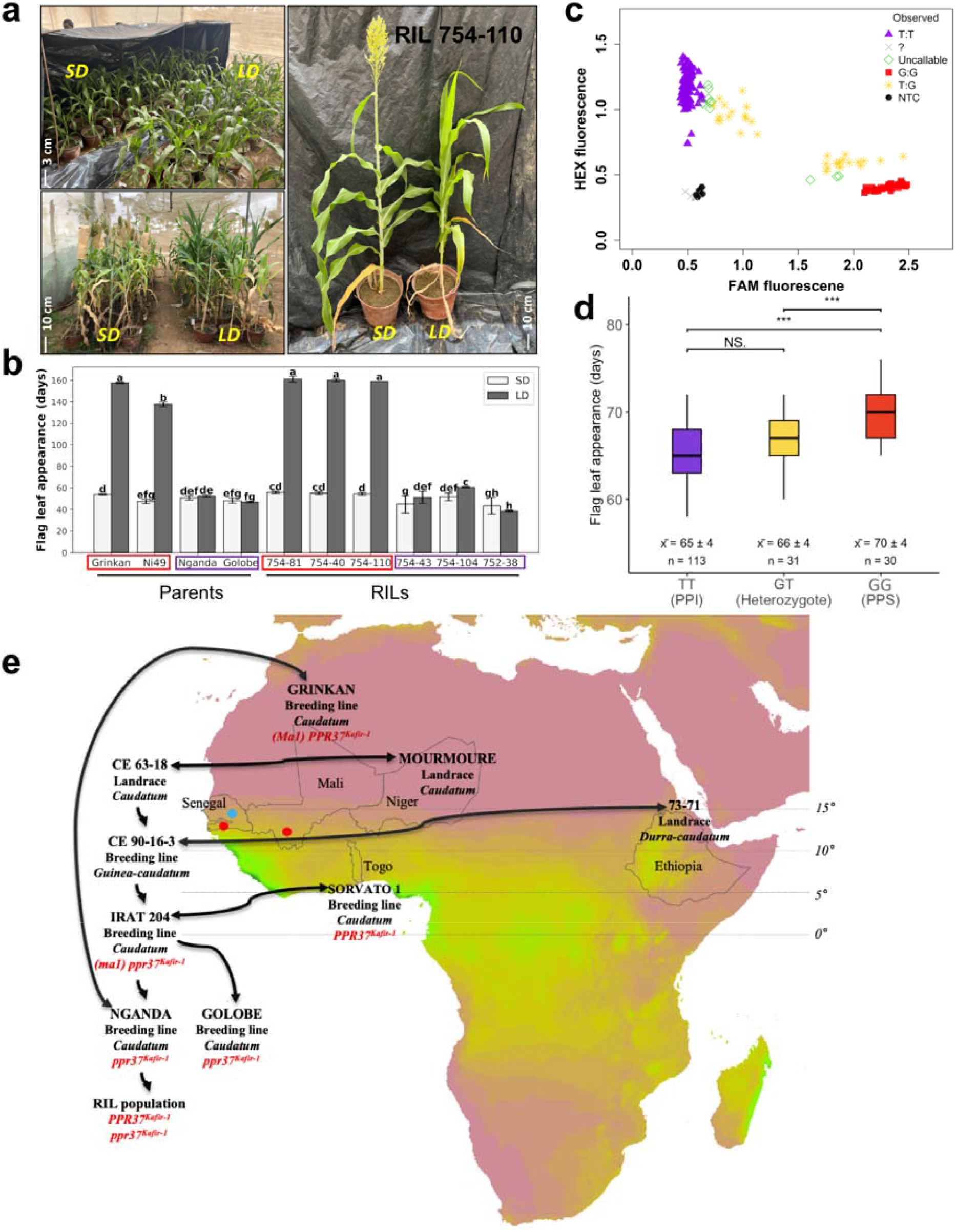
Effects of allelic variation at *Ma1* on photoperiodic flowering. (a) Experimental validation of *Ma1* allelic effect using selected photoperiod sensitive (PPS) lines carrying the *Ma1* functional allele and photoperiod-insensitive (PPI) lines carrying *ma1* loss-of function allele under short days (SD) and long days (LD) conditions. (b) Phenotypic comparison of plants between SD and LD conditions, regarding panicle initiation. Grinkan and Ni49 are photoperiod sensitive varieties used as positive controls, whereas Golobe and Nganda are photoperiod-insensitive varieties used as negative controls. Mean values with the same letters are not significantly different from one another at *p*□< □0.05 based on Least Squares Means. Red rectangles indicate PPS lines carrying the *Ma1* functional *PPR37*^*Kafir-1*^ allele and purple rectangles indicate PPI lines carrying *ma1* loss-of-function *ppr37*^*Kafir-1*^ allele. (c) Cluster plot of KASP genotyping gel-free fluorescence signal for allelic separation of the marker Sbv3.1_06_40312464K developed based on one of the causative variants at *Ma1* gene (*ppr37*^*Kafir-1*^ loss-of-function allele) in 174 F_3_ three-way cross individuals. The F_3_ population was developed based on three-way crosses, including the PPS RILs in (b), used as parents. (d) Effect of *Ma1* alleles on the F_3_ population based on marker Sbv3.1_06_40312464K, where ‘Red’ and ‘purple’ represent groups with or without the photoperiod associated allele *PPR37*^*Kafir-1*^, respectively. Yellow represents heterozygotes at the locus. NTC, no DNA-template control. Turkey test; *\*\*\*p < 0.0001; **p < 0.001; ns, not significant*. (e) Origin of the loss-of-function photoperiod-insensitive *ppr37*^*Kafir-1*^ allele (*ma1*) in the Senegalese sorghum breeding program. Double arrows indicate crosses between lines or cultivars. The background color scale on the Africa map represents the annual precipitation sum (in millimeters) from WorldClim data (Fick & Hijmans, 2017). Red and blue dots indicate the main cultivation zones of the RIL founder lines, Grinkan (800–1000 mm, in Mali with corresponding zone in Senegal) and Nganda (600–800 mm in Senegal), corresponding to the Soudanian and Soudano-Sahelian agro-climatic zones, respectively.

## 4. DISCUSSION

### 4.1. Genotype and environment interplays on photoperiodic flowering

Plant performance in the field emerges from the interplay between genotype, environment, and genotype-environment interactions. Sorghum is a widely cultivated crop adapted to diverse environmental conditions (Mace et al., 2013); however, negative effects of abiotic and biotic stresses contribute to reduced productivity and grain quality (Audilakshmi et al., 2011; Haussmann et al., 2012). In this study, we have quantified, using recombinant inbred lines, the level of GxE and the resulting photoperiodic flowering plasticity through multi-environment trials, representing the key sorghum production environments, with spillovers to their corresponding climates in Sub-Sahara Africa. The subsequent variability observed for flag leaf appearance (Figure 1b), highlights the advantage for using the RIL population as a valuable resource for dissecting GxE (Diouf & Pascual, 2021). Among various models suitable to study GxE and plasticity, the joint regression analysis enables to simultaneously assess genetically specific performance and line responses along environmental gradients, facilitating the identification of superior lines for specific environmental conditions (Lian & De Los Campos, 2016; Malosetti et al., 2016; Van Eeuwijk et al., 2016) along with their underlying genetic basis of adaptation. Our findings contribute to the overall development goal of deploying trait technology packages and understanding of adaptive trait genetic architecture to building effective elite breeding gene pools for genetic gain.

By investigating the relationship between the environmental mean and environmental parameters for floral induction with varying growth windows, we identified the diurnal temperature range within the period from 47 to 59 DAP as the critical environmental index. Early studies revealed flowering time is regulated in a complex manner by plant development (autonomous pathway), day length (photoperiod), temperature, and other factors (Grant et al., 2023; Murphy et al., 2014; Nuñez & Yamada, 2017). In the present study, the difference between the maximum temperature (day temperature) and the minimum temperature (night temperature) had a stronger effect on the variation in flag leaf appearance. This suggested that temperature may have influenced more genes involved in floral initiation than other environmental parameters (Thingnaes et al., 2003). Our findings align with a previous study, in which Mu et al. (2022) used a RIL sorghum population and found that DTR was associated with plant height, a highly heritable similar to flowering time, between 40 and 53 DAP. Both studies confirm the effect of temperature variation on flowering time as stated earlier by Myster & Moe (1995). Since flowering time is positively correlated with stem elongation in sorghum (Grant et al., 2023), both traits may be influenced by the same environmental cues, thus necessitating further investigations.

Our sorghum RIL population segregated for photoperiodism, and all six environments were considered relatively under long day condition during the growing seasons (11.4 - 12.8h) > 12 h (Lee et al., 1998). We initially hypothesized photoperiod (day length) as the key environmental parameter influencing flag leaf appearance. Instead, the correlation between photoperiod and the environmental mean was low (0.44) in the same 47–59 DAP window and increased only toward the end of trait establishment (Figure 2a). In maize, a sorghum’s close relative, Adak et al. (2024) showed that environmental cues, such as day length and radiation express their strongest effects during specific pre-flowering developmental windows. These findings, altogether, highlight the nuanced interplay and complex influence of exogenous factors on floral induction. Temperature variation on various plant responses was introduced by Went (1953), and that effect on flowering time and the effect of genes in the photoperiodic response pathway have been documented in earlier studies (Vergara & Chang, 1985). In the present study, the Grinkan allele of maturity gene *Ma1* delayed flowering in most studied environments, which supports the function of *Ma1* as a repressor of flowering under long day conditions, where the effect was dependent on temperature variation, as previously demonstrated in temperate growing environments (Murphy et al. 2014).

### 4.2. Genetic control evidence of photoperiodic plasticity for yield stability

A key hypothesis that motivated our study was that, at least, one of the known maturity loci (*Ma1-6*) (Grant et al., 2023) underlies photoperiodic plasticity along the north-south precipitation gradient in response to environmental variations as an indication of genotype adaptability. Indeed, a significant phenotypic correlation may result from the action of pleiotropic genes, which may control plasticity parameters with common loci contributing to both the phenotype baseline mean (intercept) and the plasticity response (slope), or distinct loci controlling each variable (Fu & Wang, 2023). The correlations between the flag leaf appearance mean values and plasticity measures (intercept and slope) were highly significant for both types of variables (Figure 2e). The identified common QTLs between the trait means and plasticity measures reflected these correlations at genetic level, where five of the six identified QTLs were observed for plasticity parameters (Figure S1, Table 2).

The good number of common loci were identified in our study, with a few loci having major effects (*Ma1* or *qFLA6.1, qFLA6.2*, and *qFLA6.3*) relative to the number of several independent minor effect loci (Table 2, Table S5), coupled with significant interactions among major effect loci (Table S6). These results reflect the plausible allelic sensitivity for the genetic control of photoperiodic flowering plasticity in the germplasm. The dependence between phenotype means and plasticity has been tested using tomato MAGIC and Maize NAM mapping populations, where Diouf et al. (2020) and Li et al. (2019) reported similar findings, respectively with different populations and environments. The genetic control of phenotypic plasticity has been either attributed to heterozygosity at prominent genes (overdominance), differential environmental sensitivity of alleles of a gene (allelic sensitivity), or genes that affect the expression of the phenotype and genes that react to the environment (regulatory gene model) (Ungerer et al., 2003; Via et al., 1995). On one hand, the putative allelic sensitivity found in this study, implies a trade-off for line selection because there are limited potential advantages to exploit plasticity responses while increasing phenotype mean values of genotypes (Kusmec et al., 2017). On the other hand, studies in maize have demonstrated that hybrid genotypes with higher performance were subject to the concordant phenotypic plasticity from their two founder parents (Fu & Wang, 2023), suggesting a greater performance of selected hybrid genotypes as robust and effective ways to sustain good yield performance across diverse and harsh environments.

The timing of flag leaf appearance was critical for grain mold damage at panicle maturity. In an earlier study, Thakur et al. (2007) suggested using photoperiod-sensitive varieties to avoid grain mold after finding that photoperiod-sensitive varieties could avoid grain mold by synchronizing their grain filling and maturity phase with the end of rainy seasons. The medium FLA classes (hybrid genotypes) at top two associated loci contributed more to grain mold avoidance (both PGMR and TGMR), followed by the late class at the canonical *Ma1* (*qFLA6.1*), and the early class at *qFLA6.2* in two contrasting environments, respectively, coupled with significant interactions among alleles at these top two loci (Figure 3c, Table 3, Table S6). Indeed, photoperiod-sensitive varieties can adjust their vegetative cycle to varying sowing dates and environments, thereby maintaining flowering and maturity times close to the optimal dates for high grain quality, while minimizing threats, such as bird and pathogen damage (Haussmann et al., 2012). This observation supports the concordant phenotypic plasticity model characterized in progenies co-segregating at major effect loci for photoperiodism to adapt diverse environments.

At *qFLA6.2*, genotypes with late flag leaf appearance were more affected by grain mold damage than genotypes with the medium and earlier flag leaf appearance (Table 3). This observation may challenge the notion that late flowering contributes to grain mold avoidance in sub-humid environments. In addition, *qFLA6.2* was found to be stable in environments with the highest rainfall pattern (SD), where the late FLA associated allele contributed significantly to grain yield reduction (Table 3). Negative correlations between grain mold damage and the timing of flowering have been documented previously (Audilakshmi et al., 1999; Thakur et al., 2007); here, we observed late flowering associated with *qFLA6.2*, not only contributing to greater grain mold damage but also greater grain yield reduction. This observation reveals the complexity of the disease and may indicate that fungi have different windows for maximum infection during grain development stages, on one hand. On the other hand, the effect of *qFLA6.2* may be suppressed by the presence of *Ma1* dominant allele under a more humid environment, thus leading to stable performance of hybrids genotypes with the two loci, or the plausible interactions with other unknown genetic factors at specific growth windows.

The functional causal variant at *Ma1* was the most significantly associated locus with phenotypic mean variation, the environmental slope, late maturity, grain mold, and grain yield in both the RIL and three-way cross populations under LD conditions in semi-arid (Bambey) and humid (Sedhiou) environments (Figure S1; Figure 3a; Figure 4b,d; Figure 5a,b,d; Table 3). This implies that *Ma1* is more sensitive locus for adaptation to temperature variation across environments, highlighting its potential role as adaptive response for sorghum genotypes across the Sahelian (Ethiopia to center-north of Senegal), Soudanian (center-south of Senegal and Mali) and Guinea (south most of Senegal and Togo) agro-climates (Figure 5e) and more broadly tropical climates (Klein et al., 2015). This plasticity effect of *Ma1* also suggests its potential role in small to medium scale breeding programs, particularly for improving crop resilience to harsh environmental fluctuations (Kusmec et al., 2017), where the locus have been integrated into key locally improved varieties of Senegal, Mali, and Togo following a probable past introduction from the durra-caudatum landrace, 73-71 (IS 12610) from Ethiopia in the 1980s (Figure 5e). Sequencing of *Ma1* locus across founder breeding lines from these programs, including 73-71 genotype would shed more light on the precise origin of Ma1 adaptive allele. However, the use of *Ma1* in sorghum breeding for grain mold resistance and yield stability might vary with end-use preferences. We can speculate that the functional *Ma1* allele may enhance grain quality and yield stability in regions prone to wide temperature variation, whereas it negatively impacts these traits under optimum growing conditions.

Through reaction norms definition and QTL mapping, we identified specific and plastic loci contributing to flag leaf appearance variation, and grain mold reduction along the north-south precipitation gradient in the Sahelian and Soudanian agro-climates. The allelic variant (*SbPRR37-2*) of maturity gene *Ma1* was found to be temperature-dependent in all studied environments, suggesting its adaptive role in sorghum breeding programs across Africa (Figure 5e). The strong correlation between different flag leaf appearance times and grain mold damage suggests the use of photoperiod-sensitivity to enhance sorghum grain mold avoidance and increase phenotypic plasticity. Although the same QTLs are consistently detected in different environments or parameters, the genomic intervals were relatively large for some QTLs, likely due to the limited marker density available in the sorghum DArTag mid-density SNP panel. The low marker density co-segregating in the RIL population may constitute a limitation to identify all potential genomic variants that might contribute to the observed phenotypic variation. Integration of large genetic backgrounds, combined with good genome-wide prediction models and more characterized environments would complement the identification of more climate-resilient varieties across diverse agro-climatic zones.

Moreover, with the known underlying genetic control of photoperiodic flowering on reducing grain mold and yield loss in subhumid regions, the identified major effect maturity loci (*Ma1, qFLA6.2*, and *qFLA6.3*) would contribute to effectively building de novo elite breeding gene pools by designing iso-elite (compatible) crosses that can facilitate genetic gain (Fall et al., 2026). Overall, the selected stable genotypes and identified stable loci provide new trait technology and tools for understanding sorghum climate-resilience and building stable elite breeding for genetic gain.

## ABBREVIATIONS

BLUE: best linear unbiased estimate
BLUP: best linear unbiased predictor
CERAAS: centre d’etude régional pour l’amélioration de l’adaptation à la sécheresse
CERIS: critical environmental regressor through the informed search
CIM: composite interval mapping
CNRA: centre national de recherches agronomiques
DL: daylength
DTF: days to flowering
DTR: diurnal temperature range
EMS: Ethyl Methanesulfonate
FLA: flag leaf appearance
GDD: growing degree day
GxE: genotype-environment interaction
KASP: Kompetitive allele-specific polymerase chain reaction
LD: long day
LOD: logarithm of odds
PGMR: panicle grain mold rate at physiological maturity
PRCP: precipitation
PTR: photothermal ratio
PTT: photothermal time
PVE: phenotype variation estimated
QTL: quantitative trait locus
SD: short day
SNP: single nucleotide polymorphism
TGMR: grain mold rate after threshing
YLD: grain yield

## ACKNOWLEDGMENTS

This study is made possible by the support of the American People provided to the Feed the Future Innovation Lab for Crop Improvement through the United States Agency for International Development (USAID) and by the Bill and Melinda Gates Foundation (BMGF) project subgrant G-40792-03, Green Evolution - Accelerating Dryland Cereals Improvement for Africa. The first author acknowledges the Ph.D. In-Region Scholarship provided by the German Academy Service Exchange (DAAD).

## AUTHOR CONTRIBUTIONS

CD, JMF, NAK, GPM conceived the research; JMF, DATH designed the study; DATH, SB, MN performed the experiments and genotyping; CD, SB developed mapping populations; TF, GPM developed molecular markers; CD, NAK, GPM, DD, BS, DS performed data curation, experimental design and supervised the study; NAK, CD, GPM, JMF provided funding resources; DATH performed data analysis and visualization; DATH and JMF wrote the original draft. All authors wrote, reviewed, and approved the manuscript.

## DATA AVAILABILITY STATEMENT

The phenotypic, genotypic data, and R scripts will be available at Dryad Digital Repository (DOI: 10.5061/dryad.z8w9ghxs1) following acceptance.

## Supplemental Materials

**Figure S1:**
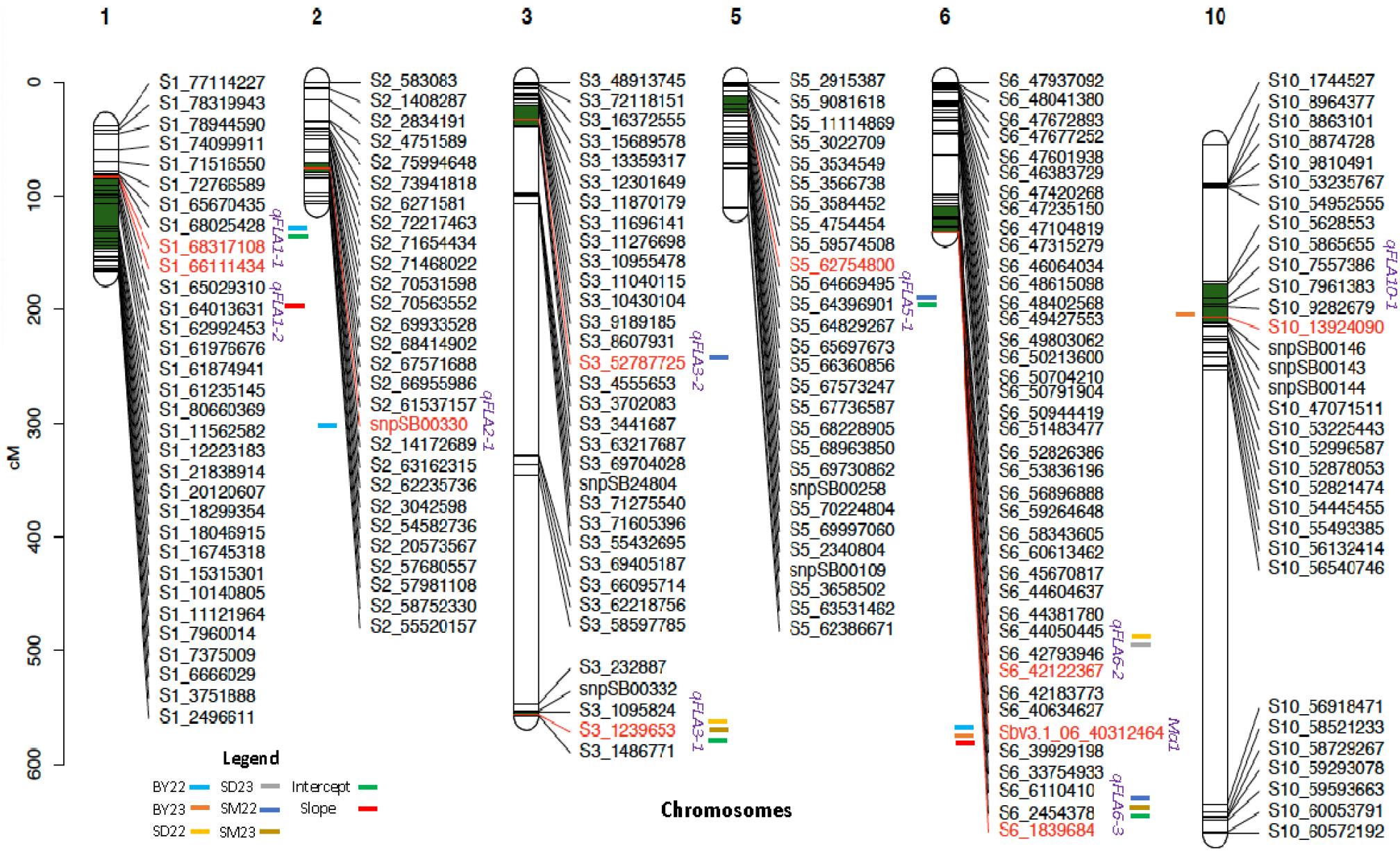
Genetic positions of the significantly associated QTLs for flag leaf appearance. The single nucleotide polymorphism (SNP) markers representing the QTL are highlighted in red on each chromosome. A LOD (logarithm of the odds) significance threshold of 5.9 was used to declare the association of a genomic region with the trait variation. The position of each QTL confidence interval is indicated by green bars on the chromosome. The color of the horizontal bar next to the QTL or marker name indicates the environment and/or reaction-norm parameter (intercept or slope) for which the QTL was detected using FLA values from three contrasting locations, Bambey (BY), Sinthiou Maleme (SM) and Sedhiou (SD) over two years (2022 and 2023). Marker names starting with the string, snpSB or Sbv are trait linked KASP markers present in the sorghum mid-density genotyping marker panel (DArTag panel).

**Figure S2:**
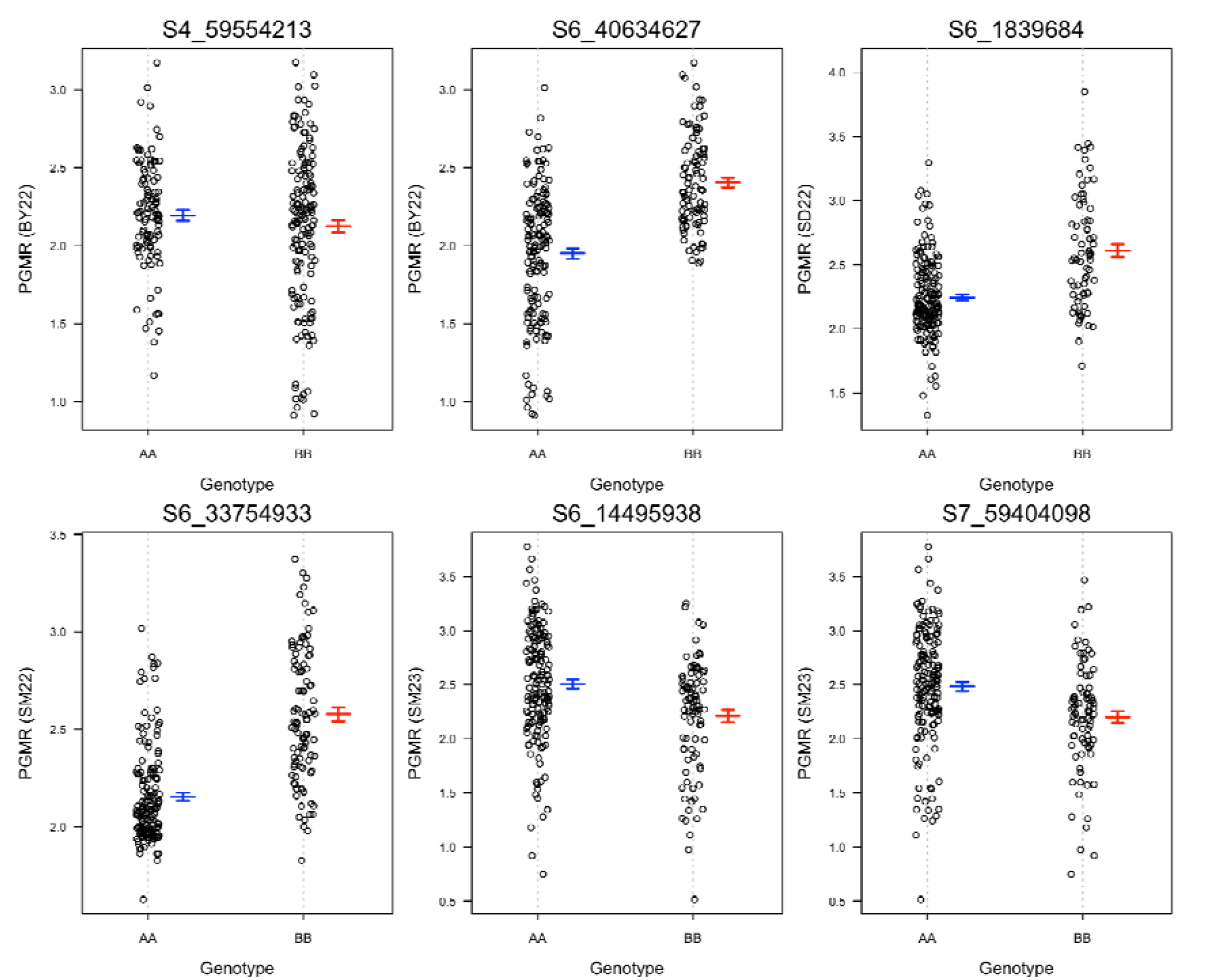
Genotype–phenotype plots for significant QTL associated with grain mold damage at panicle maturity (PGMR). AA = Grinkan, BB = Nganda.

**Figure S3:**
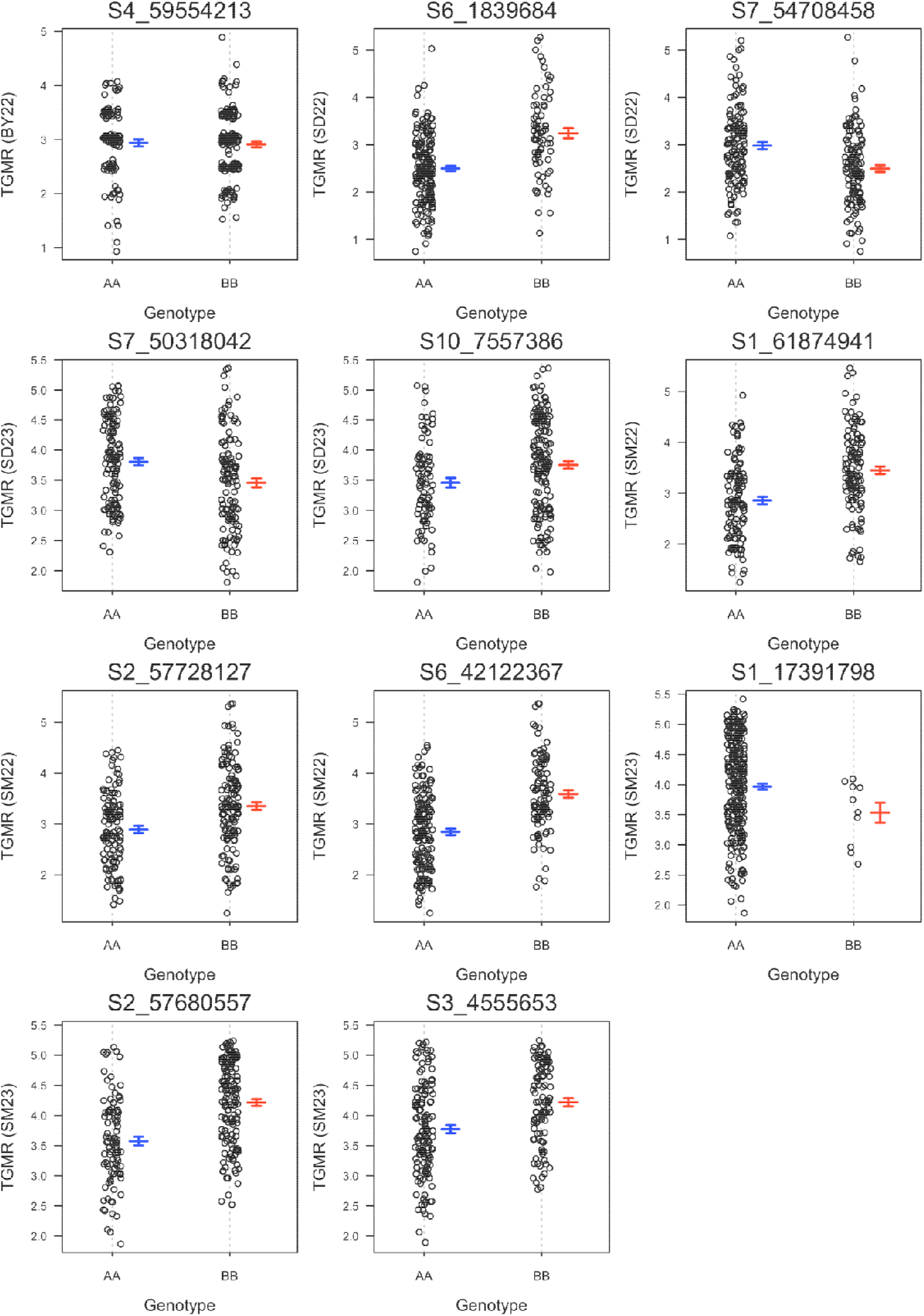
Genotype–phenotype plots for significant QTL associated with threshed grain mold damage (TGMR). AA = Grinkan, BB = Nganda.

**Figure S4:**
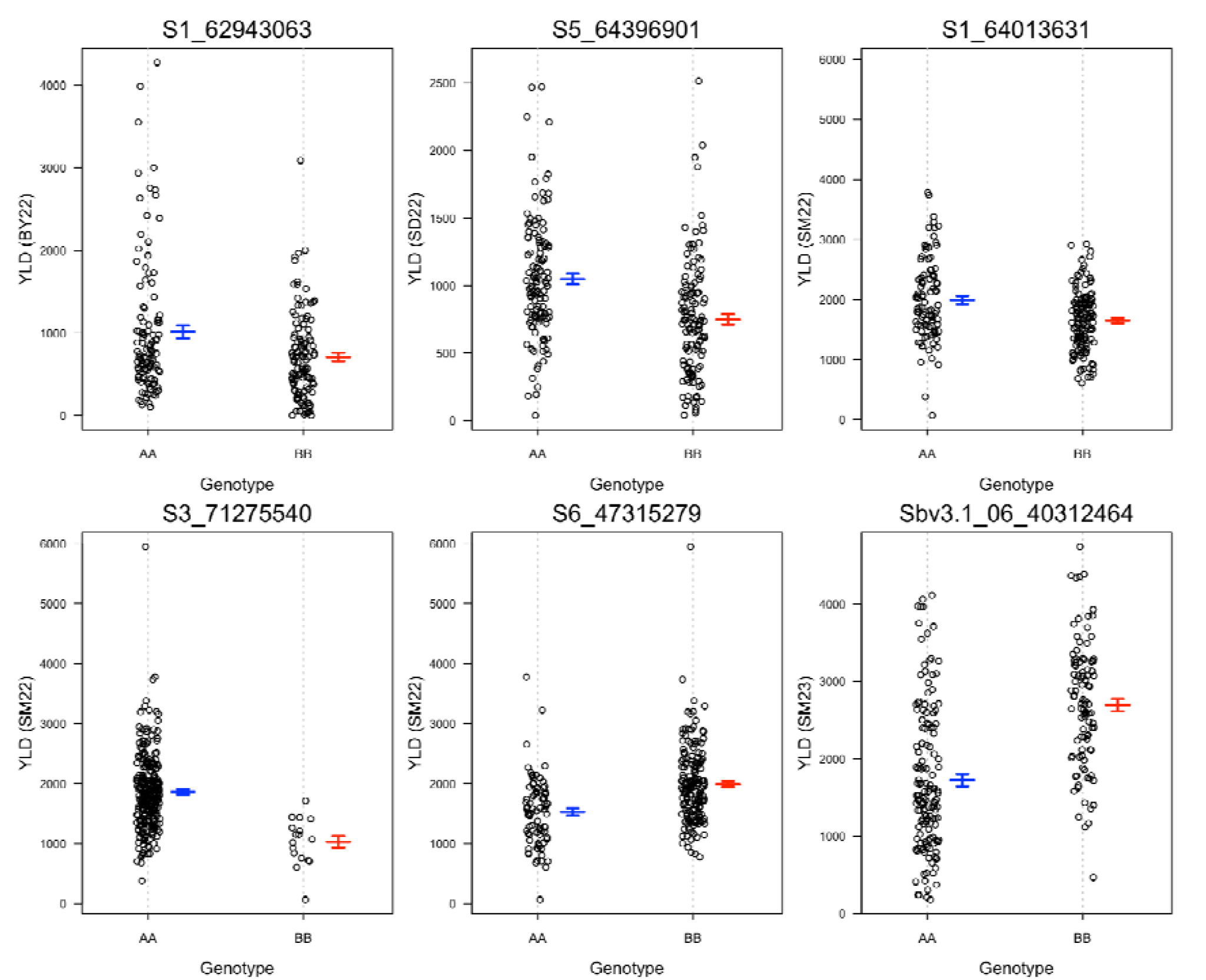
Genotype–phenotype plots for significant QTL associated with grain yield (YLD). AA = Grinkan, BB = Nganda.

**Table S1:** Information on field trials carried out in the present study.

| Location | Latitude | Year | Abbreviation | Planting date | Harvest date |
| --- | --- | --- | --- | --- | --- |
| Bambey<br>(Semi-arid) | 14°41'N | 2022 | BY22 | 2022-07-06 | 2022-11-11 |
|  |  | 2023 | BY23 | 2023-07-17 | 2023-11-17 |
| Sinthiou Maleme<br>(Sub-humid) | 13°49'N | 2022 | SM22 | 2022-07-29 | 2022-12-08 |
|  |  | 2023 | SM23 | 2023-07-19 | 2023-11-22 |
| Sedhiou<br>(Humid) | 12°49'N | 2022 | SD22 | 2022-07-19 | 2022-11-30 |
|  |  | 2023 | SD23 | 2023-07-20 | 2023-11-24 |

**Table S2:**
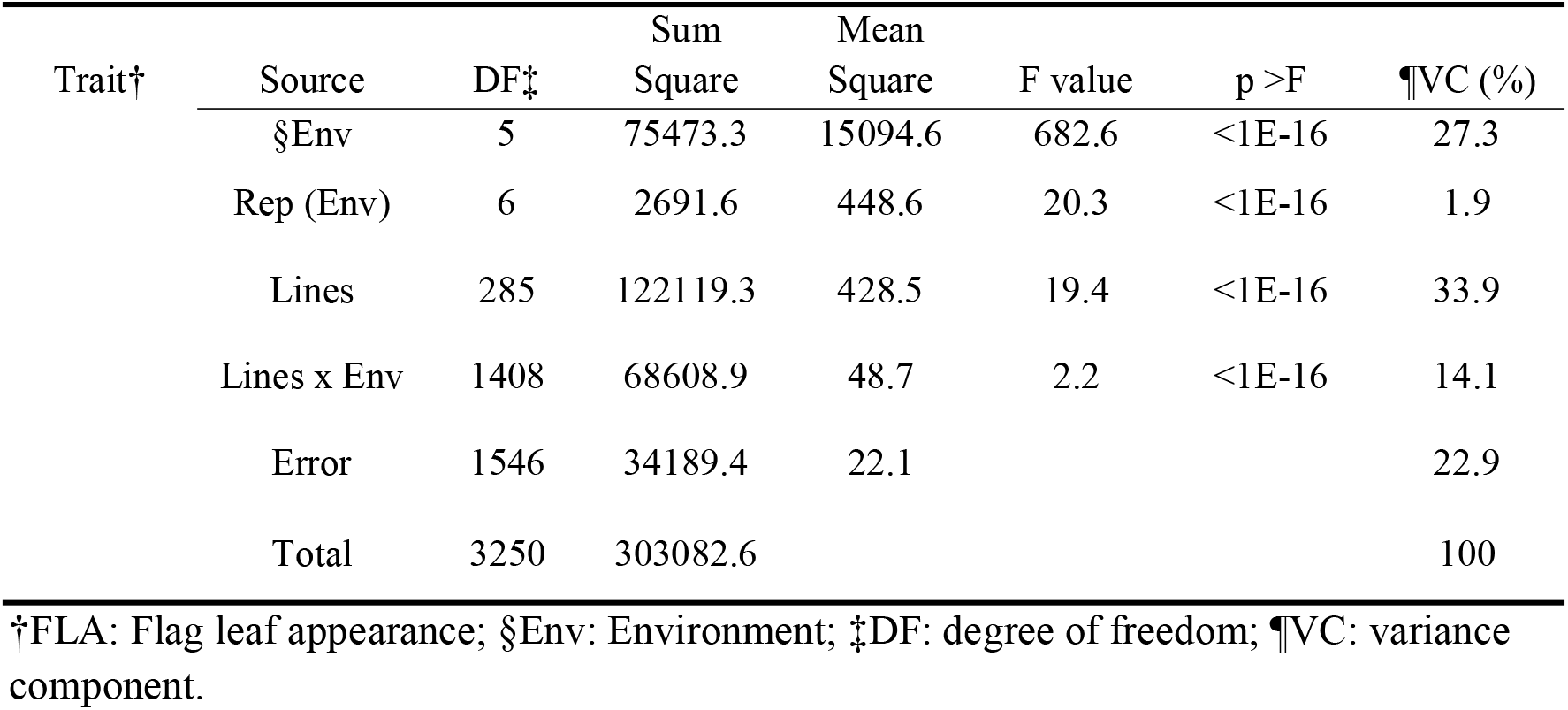
Summary statistics of analysis of variance for FLA from the combined analysis of six environments.

**Table S3:** Analysis of variance for flag leaf appearance following the regression on the mean model (Finlay-Wilkinson model).

| Trait | Source | DF | Sum Square | Mean Square | F value | p > F |
| --- | --- | --- | --- | --- | --- | --- |
| FLA | Environments | 5 | 36823 | 7364.8 | 264.9 | <0.001 |
|  | Genotype | 285 | 62426 | 219 | 7.9 | <0.001 |
|  | Heterogeneity of slopes | 285 | 3171 | 11.1 | 0.4 | <0.001 |
|  | Error | 1140 | 31700 | 27.8 | - |  |
|  | Total | 1715 | 134121 | - | - | - |

FLA: Flag leaf appearance; DF: degree of freedom.

**Table S4:**
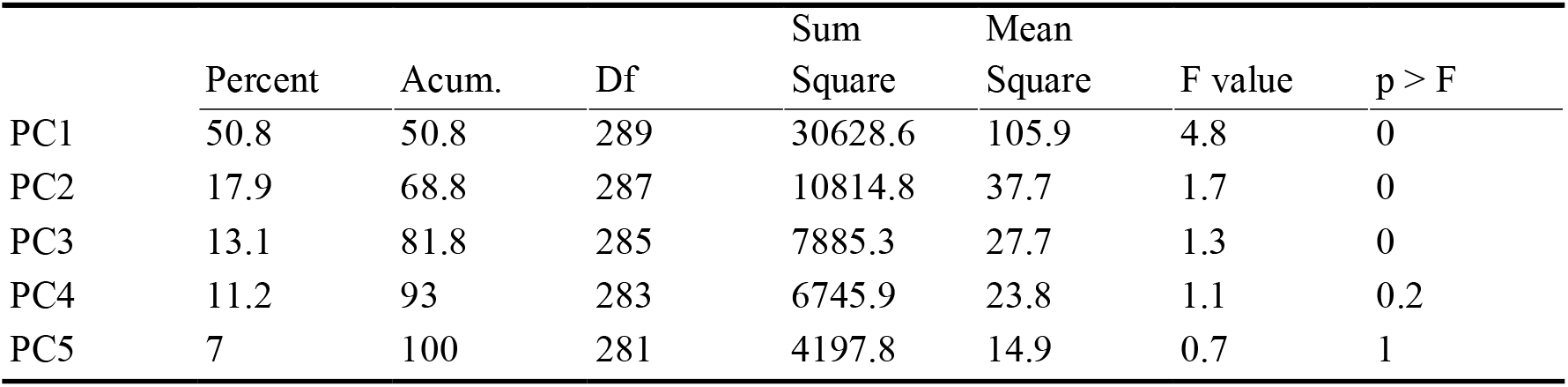
Analysis of variance for flag leaf appearance following the regression on the mean model (AMMI model).

|  | Percent | Acum. | Df | Sum<br>Square | Mean<br>Square | F value | p > F |
| --- | --- | --- | --- | --- | --- | --- | --- |
| PC1 | 50.8 | 50.8 | 289 | 30628.6 | 105.9 | 4.8 | 0 |
| PC2 | 17.9 | 68.8 | 287 | 10814.8 | 37.7 | 1.7 | 0 |
| PC3 | 13.1 | 81.8 | 285 | 7885.3 | 27.7 | 1.3 | 0 |
| PC4 | 11.2 | 93 | 283 | 6745.9 | 23.8 | 1.1 | 0.2 |
| PC5 | 7 | 100 | 281 | 4197.8 | 14.9 | 0.7 | 1 |

**Table S5:**
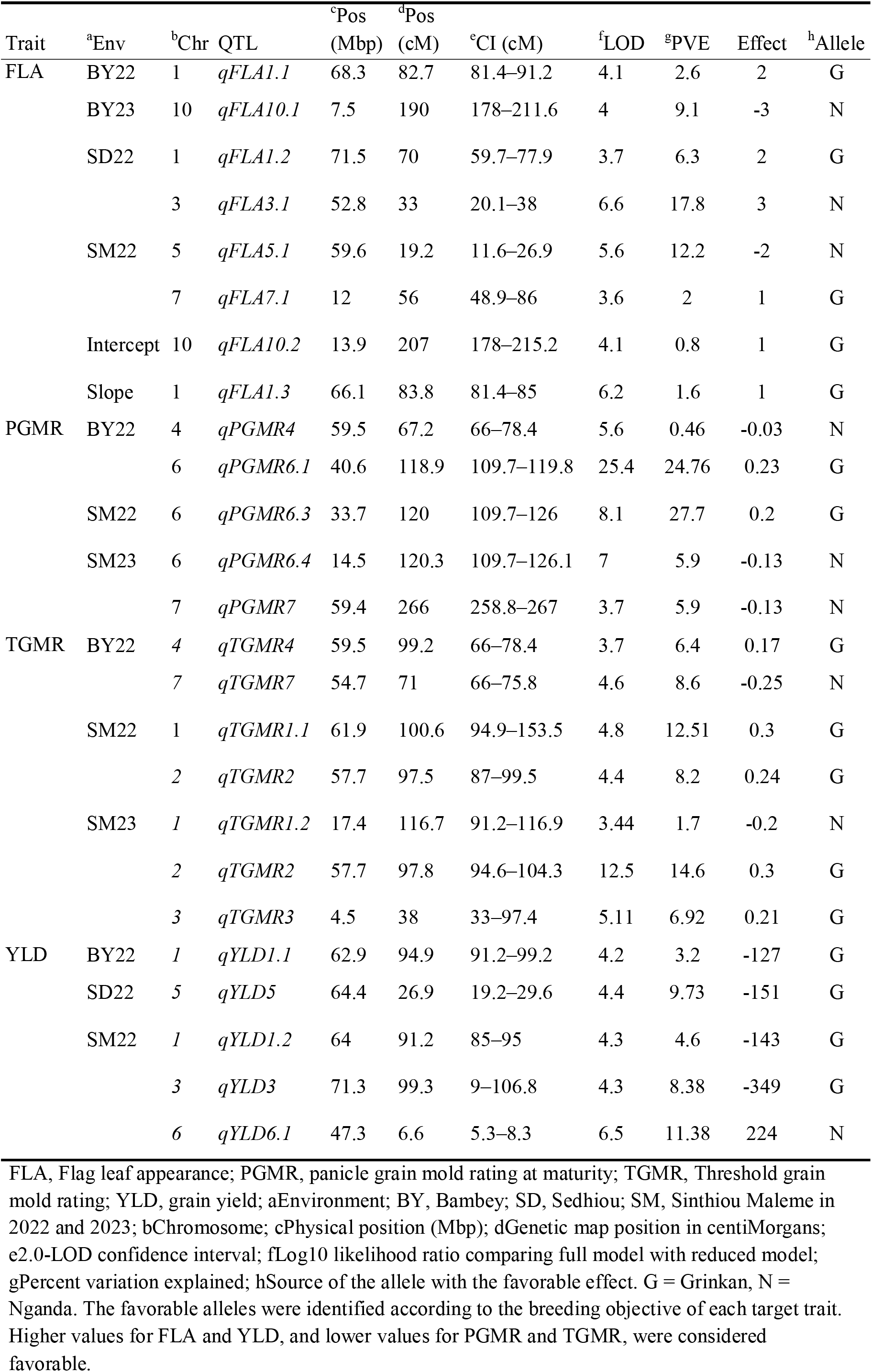
The top non-overlapping QTLs identified for FLA, plasticity measures of FLA, PGMR, TGMR, and YLD in contrasting environments.

**Table S6:**
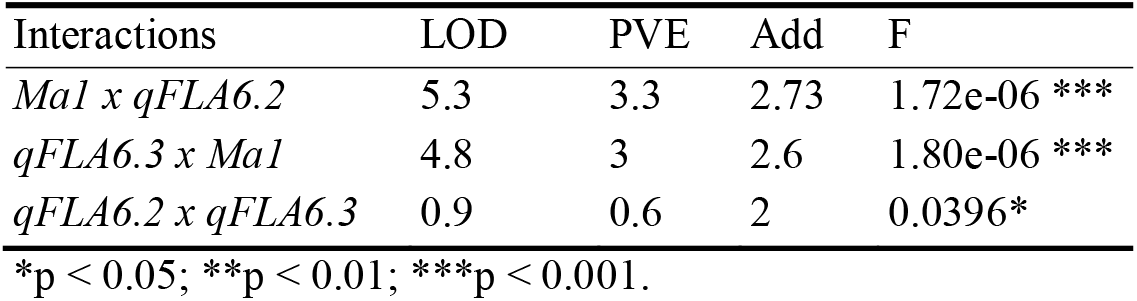
Interactions among stable QTLs on chromosome 6.

## REFERENCE

Adak, A., Murray, S. C., Varela, J. I., Infante, V., Wilker, J., Calderón, C. I., Subramanian, N., de Leon, N., Yu, J., Stull, M. A., Brun, M., Hill, J., Johnson, C. D., Riera-Lizarazu, O., Rooney, W. L., & Zhang, H. (2024). Photoperiod associated late flowering reaction norm: Dissecting loci and genomic-enviromic associated prediction in maize. Field Crops Research, 311, 109380. 10.1016/j.fcr.2024.109380

Ambekar, S. S., Kamatar, M. Y., Ganesamurthy, K., Ghorade, R. B., Saxena, U., Chand, P., Jadav, B. D., Das, I. K., Nageshwararao, T. G., Audilakshmi, S., & Seetharama, N. (2011). Genetic enhancement of sorghum (*Sorghum bicolor* (L) Moench) for grain mould resistance: II. Breeding for grain mould resistance. Crop Protection, 30(7), 759–764. 10.1016/j.cropro.2010.06.024

Aruna, C., Audilakshmi, S., Ratnavathi, C. v., & Patil, J. v. (2012). Grain quality improvement of rainy season sorghums. Quality Assurance and Safety of Crops & Foods, 4(3), 149–149. 10.1111/j.1757-837X.2012.00157.x

Aruna, C., Das, I. K., Reddy, P. S., Ghorade, R. B., Gulhane, A. R., Kalpande, V. V., Kajjidoni, S. T., Hanamaratti, N. G., Chattannavar, S. N., Mehtre, S., Gholve, V., Kamble, K. R., Deepika, C., Kannababu, N., Bahadure, D. M., Govindaraj, M., & Tonapi, V. A. (2021). Development of sorghum genotypes for improved yield and resistance to grain mold using population breeding approach. Frontiers in Plant Science, 12. 10.3389/fpls.2021.687332

Audilakshmi, S., Das, I. K., Ghorade, R. B., Mane, P. N., Kamatar, M. Y., Narayana, Y. D., & Seetharama, N. (2011). Genetic improvement of sorghum for grain mould resistance: I. Performance of sorghum recombinant inbred lines for grain mould reactions across environments. Crop Protection, 30(7), 753–758. 10.1016/j.cropro.2010.12.024

Audilakshmi, S., Stenhouse, J. W., & Reddy, T. P. (2005). Genetic analysis of grain mold resistance in white seed sorghum genotypes. Euphytica, 145(1), 95–101. 10.1007/s10681-005-0534-6

Audilakshmi, S., Stenhouse, J. W., Reddy, T. P., & Prasad, M. V. R. (1999). Grain mould resistance and associated characters of sorghum genotypes. Euphytica, 107(2), 91–103. 10.1023/A:1026410913896

Bandyopadhyay, R., Butler, D. R., Chandrashekar, A., Reddy, R. K., & Navi, S. S. (2000). Biology, epidemiology, and management of sorghum grain mold. Technical and Institutional Options for Sorghum Grain Mold Management: Proceedings of an International Consultation, 18–19.

Bates, D., Mächler, M., Bolker, B., & Walker, S. (2015). Fitting linear mixed-effects models using lme4. Journal of Statistical Software, 67, 1–48. 10.18637/jss.v067.i01

Bhosale, S. U., Stich, B., Rattunde, H. F. W., Weltzien, E., Haussmann, B. I., Hash, C. T., Ramu, P., Cuevas, H. E., Paterson, A. H., Melchinger, A. E., & Parzies, H. K. (2012). Association analysis of photoperiodic flowering time genes in west and central African sorghum [Sorghum bicolor (L.) Moench]. BMC Plant Biology, 12(1), 32. 10.1186/1471-2229-12-32

Borrell, A., van Oosterom, E., George-Jaeggli, B., Vadez, V., Singh, V., & Hammer, G. (2020). Physiology of growth, development and yield. In V. A. Tonapi, H. S. Talwar, A. K. Are, B. V. Bhat, Ch. R. Reddy, & T. J. Dalton (Eds.), Sorghum in the 21st Century: Food – Fodder – Feed – Fuel for a Rapidly Changing World (pp. 127–155). Springer. 10.1007/978-981-15-8249-3_6

Broman, K. W., Wu, H., Sen, Ś., & Churchill, G. A. (2003). R/qtl: QTL mapping in experimental crosses. Bioinformatics, 19(7), 889–890. 10.1093/bioinformatics/btg112

Casto, A. L., Mattison, A. J., Olson, S. N., Thakran, M., Rooney, W. L., & Mullet, J. E. (2019). Maturity2, a novel regulator of flowering time in sorghum bicolor, increases expression of SbPRR37 and SbCO in long days delaying flowering. PLOS ONE, 14(4), e0212154. 10.1371/journal.pone.0212154

Chen, Y., Xiong, Y., Hong, H., Li, G., Gao, J., Guo, Q., Sun, R., Ren, H., Zhang, F., Wang, J., Song, J., & Qiu, L. (2023). Genetic dissection of and genomic selection for seed weight, pod length, and pod width in soybean. The Crop Journal, 11(3), 832–841. 10.1016/j.cj.2022.11.006

Costa-Neto, G., Galli, G., Carvalho, H. F., Crossa, J., & Fritsche-Neto, R. (2021). EnvRtype: A software to interplay enviromics and quantitative genomics in agriculture. G3 Genes|Genomes|Genetics, 11(4), jkab040. 10.1093/g3journal/jkab040

Cuevas, H. E., Fermin-Pérez, R. A., Prom, L. K., Cooper, E. A., Bean, S., & Rooney, W. L. (2019). Genome-wide association mapping of grain mold resistance in the US sorghum association panel. The Plant Genome, 12(2), 180070. 10.3835/plantgenome2018.09.0070

Dalton, T. J., & Hodjo, M. (2020). Trends in global production, consumption, and utilization of sorghum. In V. A. Tonapi, H. S. Talwar, A. K. Are, B. V. Bhat, Ch. R. Reddy, & T. J. Dalton (Eds.), Sorghum in the 21st Century: Food – Fodder – Feed – Fuel for a Rapidly Changing World (pp. 3–15). Springer. 10.1007/978-981-15-8249-3_1

Danecek, P., Auton, A., Abecasis, G., Albers, C. A., Banks, E., DePristo, M. A., Handsaker, R. E., Lunter, G., Marth, G. T., Sherry, S. T., McVean, G., Durbin, R., & 1000 Genomes Project Analysis Group. (2011). The variant call format and VCFtools. Bioinformatics, 27(15), 2156–2158. 10.1093/bioinformatics/btr330

Das, I., Aruna, C., & Tonapi, V. (2020). Sorghum grain mold. ICAR-Indian Institute of Millets Research, Hyderabad, India, 1–86.

Das, I., Audilakshmi, S., & Patil, J. (2012). Fusarium grain mold: The major component of grain mold disease complex in sorghum (Sorghum bicolor L. Moench). Eur. J. Plant Sci. Biotechnol, 6, 45–55.

Diatta, C., Sarr, M. P., Tovignan, T. K., Aidara, O., Dzidzienyo, D. K., Diatta-Holgate, E., Faye, J. M., Danquah, E. Y., Offei, S. K., & Cisse, N. (2021). Multienvironment evaluation of tannin-free photoperiod-insensitive sorghum (*Sorghum bicolor* (L) Moench) for yield and resistance to grain mold in Senegal. International Journal of Agronomy, 2021, e5534314. 10.1155/2021/5534314

Diatta, C., Tovignan, T. K., Adoukonou-Sagbadja, H., Aidara, O., Diao, Y., Sarr, M. P., Ifie, B. E., Offei, S. K., Danquah, E. Y., & Cisse, N. (2019). Development of sorghum hybrids for stable yield and resistance to grain mold for the Center and South-East of Senegal. Crop Protection, 119, 197–207. 10.1016/j.cropro.2019.02.001

Dingkuhn, M., Kouressy, M., Vaksmann, M., Clerget, B., & Chantereau, J. (2008). A model of sorghum photoperiodism using the concept of threshold-lowering during prolonged appetence. European Journal of Agronomy, 28(2), 74–89. 10.1016/j.eja.2007.05.005

Diouf, I., Derivot, L., Koussevitzky, S., Carretero, Y., Bitton, F., Moreau, L., & Causse, M. (2020). Genetic basis of phenotypic plasticity and genotype × environment interactions in a multi-parental tomato population. Journal of Experimental Botany, 71(18), 5365–5376. 10.1093/jxb/eraa265

Diouf, I., & Pascual, L. (2021). Multiparental population in crops: Methods of development and dissection of genetic traits. In P. Tripodi (Ed.), Crop Breeding (Vol. 2264, pp. 13–32). Springer US. 10.1007/978-1-0716-1201-9_2

Excellence in breeding. (2022). CGIAR Genetic Innovation (GI) Toolbox. Excellence in Breeding. https://excellenceinbreeding.org/toolbox/services/middensity-genotyping-service

Fall, S. T., Kena, A., Rice, B. R., Kanfany, G., Diatta, C., Kane, N. A., Fritz, A. K., & Morris, G. P. (2026). Genomic approaches to build de novo elite breeding gene pools from locally adapted landraces. Theoretical and Applied Genetics, 139(1), 28. 10.1007/s00122-025-05124-2

FAO. (2024). FAOSTAT database. https://www.fao.org/faostat/fr/#home

Fu, R., & Wang, X. (2023). Modeling the influence of phenotypic plasticity on maize hybrid performance. Plant Communications, 4(3), 100548. 10.1016/j.xplc.2023.100548

Gauch, H. G., & Zobel, R. W. (1990). Imputing missing yield trial data. Theoretical and Applied Genetics, 79(6), 753–761. 10.1007/BF00224240

Gollob, H. F. (1968). A statistical model which combines features of factor analytic and analysis of variance techniques. Psychometrika, 33(1), 73–115.

Grant, N. P., Toy, J. J., Funnell-Harris, D. L., & Sattler, S. E. (2023). Deleterious mutations predicted in the sorghum (Sorghum bicolor) Maturity (Ma) and Dwarf (Dw) genes from whole-genome resequencing. Scientific Reports, 13(1), 16638. 10.1038/s41598-023-42306-8

Guitton, B., Théra, K., Tékété, M. L., Pot, D., Kouressy, M., Témé, N., Rami, J.-F., & Vaksmann, M. (2018). Integrating genetic analysis and crop modeling: A major QTL can finely adjust photoperiod-sensitive sorghum flowering. Field Crops Research, 221, 7–18. 10.1016/j.fcr.2018.02.007

Hardenbol, P., Banér, J., Jain, M., Nilsson, M., Namsaraev, E. A., Karlin-Neumann, G. A., Fakhrai-Rad, H., Ronaghi, M., Willis, T. D., Landegren, U., & Davis, R. W. (2003). Multiplexed genotyping with sequence-tagged molecular inversion probes. Nature Biotechnology, 21(6), 673–678. 10.1038/nbt821

Harlan, J. R., & de Wet, J. M. J. (1972). A Simplified classification of cultivated sorghum. Crop Science, 12(2). 10.2135/cropsci1972.0011183X001200020005x

Haussmann, B. I. G., Fred Rattunde, H., Weltzien-Rattunde, E., Traoré, P. S. C., vom Brocke, K., & Parzies, H. K. (2012). Breeding strategies for adaptation of pearl millet and sorghum to climate variability and change in West Africa. Journal of Agronomy and Crop Science, 198(5), 327–339. 10.1111/j.1439-037X.2012.00526.x

Holland, J. B., Nyquist, W. E., & Cervantes-Martínez, C. T. (2002). Estimating and interpreting heritability for plant breeding: An update. In Plant Breeding Reviews (pp. 9–112). John Wiley & Sons, Ltd. 10.1002/9780470650202.ch2

Hossain, Md. S., Islam, Md. N., Rahman, Md. M., Mostofa, M. G., & Khan, Md. A. R. (2022). Sorghum: A prospective crop for climatic vulnerability, food and nutritional security. Journal of Agriculture and Food Research, 8, 100300. 10.1016/j.jafr.2022.100300

Kane, N. A., Foncéka, D., & Dalton, T. J. (2022). Crop adaptation and improvement for drought-Prone environments. 547-p.

Kena, A. W., Tetteh, I. T., & Burgos, C. C. (2025). panGenomeBreedr: Helpers for pangenome-enabled crop breeding. https://github.com/awkena/panGenomeBreedr

Klein, R. R., Miller, F. R., Dugas, D. V., Brown, P. J., Burrell, A. M., & Klein, P. E. (2015). Allelic variants in the PRR37 gene and the human-mediated dispersal and diversification of sorghum. Theoretical and Applied Genetics, 128(9), 1669–1683. 10.1007/s00122-015-2523-z

Klein, R. R., Rodriguez-Herrera, R., Schlueter, J. A., Klein, P. E., Yu, Z. H., & Rooney, W. L. (2001). Identification of genomic regions that affect grain-mould incidence and other traits of agronomic importance in sorghum. Theoretical and Applied Genetics, 102(2), 307–319. 10.1007/s001220051647

Kumar, R., Ikerd, J., Karthikeyan, R., & Kousik, C. (2025). Fine mapping, introgression, and KASP marker development for powdery mildew resistance in watermelon using an interspecific RIL population (Citrullus mucosospermus[×[C. lanatus). Theoretical and Applied Genetics, 138(12), 299. 10.1007/s00122-025-05079-4

Kusmec, A., Srinivasan, S., Nettleton, D., & Schnable, P. S. (2017). Distinct genetic architectures for phenotype means and plasticities in Zea mays. Nature Plants, 3(9), 715–723. 10.1038/s41477-017-0007-7

Lee, I.-J., Foster, K. R., & Morgan, P. W. (1998). Photoperiod control of gibberellin levels and flowering in sorghum. Plant Physiology, 116(3), 1003–1011. 10.1104/pp.116.3.1003

Li, C., Wu, X., Li, Y., Shi, Y., Song, Y., Zhang, D., Li, Y., & Wang, T. (2019). Genetic architecture of phenotypic means and plasticities of kernel size and weight in maize. Theoretical and Applied Genetics, 132(12), 3309–3320. 10.1007/s00122-019-03426-w

Li, X., Guo, T., Mu, Q., Li, X., & Yu, J. (2018). Genomic and environmental determinants and their interplay underlying phenotypic plasticity. Proceedings of the National Academy of Sciences, 115(26), 6679–6684. 10.1073/pnas.1718326115

Lian, L., & de los Campos, G. (2016). FW: An R package for Finlay–Wilkinson regression that incorporates genomic/pedigree information and covariance structures between environments. G3 Genes|Genomes|Genetics, 6(3), 589–597. 10.1534/g3.115.026328

Little, C. R. (2000). Plant responses to early infection events in sorghum grain mold interactions. Technical and Institutional Options for Sorghum Grain Mold Management. Proc. Int. Consultation, 18–19.

Mace, E. S., Tai, S., Gilding, E. K., Li, Y., Prentis, P. J., Bian, L., Campbell, B. C., Hu, W., Innes, D. J., Han, X., Cruickshank, A., Dai, C., Frère, C., Zhang, H., Hunt, C. H., Wang, X., Shatte, T., Wang, M., Su, Z., … Wang, J. (2013). Whole-genome sequencing reveals untapped genetic potential in Africa’s indigenous cereal crop sorghum. Nature Communications, 4(1), Article 1. 10.1038/ncomms3320

Maina, F., Faye, J. M., Kena, A. W., Diatta, C., Tankari, M. I., Ardaly, O. A., Diakite, O. S., Ibrahim, A. M., Moussa, O. A., Harou, A., & others. (2025). Delivering trait-enhanced varieties to African smallholders through a pangenomic breeding network. bioRxiv, 2025–08.

Malosetti, M., Bustos-Korts, D., Boer, M. P., & van Eeuwijk, F. A. (2016). Predicting responses in multiple environments: Issues in relation to genotype × environment interactions. Crop Science, 56(5), 2210–2222. 10.2135/cropsci2015.05.0311

Margarido, G. R. A., Souza, A. P., & Garcia, A. A. F. (2007). OneMap: Software for genetic mapping in outcrossing species: OneMap. Hereditas, 144(3), 78–79. 10.1111/j.2007.0018-0661.02000.x

Mu, Q., Guo, T., Li, X., & Yu, J. (2022). Phenotypic plasticity in plant height shaped by interaction between genetic loci and diurnal temperature range. New Phytologist, 233(4), 1768–1779. 10.1111/nph.17904

Murphy, R. L., Morishige, D. T., Brady, J. A., Rooney, W. L., Yang, S., Klein, P. E., & Mullet, J. E. (2014). Ghd7 (Ma6) represses sorghum flowering in long days: Ghd7 alleles enhance biomass accumulation and grain production. The Plant Genome, 7(2). 10.3835/plantgenome2013.11.0040

Myster, J., & Moe, R. (1995). Effect of diurnal temperature alternations on plant morphology in some greenhouse crops—A mini review. Scientia Horticulturae, 62(4), 205–215. 10.1016/0304-4238(95)00783-P

Nuñez, F. D. B., & Yamada, T. (2017). Molecular regulation of flowering time in grasses. Agronomy, 7(1), Article 1. 10.3390/agronomy7010017

Ongom, P. O., Fatokun, C., Togola, A., Garcia-Oliveira, A. L., Ng, E. H., Kilian, A., Lonardi, S., Close, T. J., & Boukar, O. (2024). A mid-density single-nucleotide polymorphism panel for molecular applications in cowpea (Vigna unguiculata (L.) Walp). International Journal of Genomics, 2024(1), 9912987. 10.1155/2024/9912987

Patterson, H. D., & Thompson, R. (1971). Recovery of inter-block information when block sizes are unequal. Biometrika, 58(3), 545–554. 10.1093/biomet/58.3.545

Placinta, C. M., D’Mello, J. P. F., & Macdonald, A. M. C. (1999). A review of worldwide contamination of cereal grains and animal feed with *Fusarium* mycotoxins. Animal Feed Science and Technology, 78(1), 21–37. 10.1016/S0377-8401(98)00278-8

R Core Team. (2025). R: A language and environment for statistical computing. R Foundation for Statistical Computing. https://www.R-project.org/

Rahman, Md. Z., Hasan, Md. T., & Rahman, J. (2023). Kompetitive Allele-Specific PCR (KASP): An efficient high-throughput genotyping platform and Its applications in crop variety development. In N. Kumar (Ed.), Molecular marker techniques: A potential approach of crop improvement (pp. 25–54). Springer Nature. 10.1007/978-981-99-1612-2_2

Rodríguez-Álvarez, M. X., Boer, M. P., van Eeuwijk, F. A., & Eilers, P. H. C. (2018). Correcting for spatial heterogeneity in plant breeding experiments with P-splines. Spatial Statistics, 23, 52–71. 10.1016/j.spasta.2017.10.003

Sparks, A. H. (2018). nasapower: A NASA power global meteorology, surface solar energy and climatology data client for R. Journal of Open Source Software, 3(30), 1035. 10.21105/joss.01035

Thakur, R. P., Reddy, B. V. S., & Mathur, K. (2007). Screening techniques for sorghum diseases. Information (Vol. 76). International Crops Research Institute for the Semi-Arid Tropics. http://oar.icrisat.org/4062/

Thingnaes, E., Torre, S., Ernstsen, A., & Moe, R. (2003). Day and night temperature responses in arabidopsis: Effects on gibberellin and auxin content, cell size, morphology and flowering time. Annals of Botany, 92(4), 601–612. 10.1093/aob/mcg176

Ungerer, M. C., Halldorsdottir, S. S., Purugganan, M. D., & Mackay, T. F. C. (2003). Genotype-environment interactions at quantitative trait loci affecting inflorescence development in arabidopsis thaliana. Genetics, 165(1), 353–365. 10.1093/genetics/165.1.353

Upadhyaya, H. D., Wang, Y.-H., Sharma, R., & Sharma, S. (2013). SNP markers linked to leaf rust and grain mold resistance in sorghum. Molecular Breeding, 32(2), 451–462. 10.1007/s11032-013-9883-3

van Eeuwijk, F. A., Bustos-Korts, D. V., & Malosetti, M. (2016). What should students in plant breeding Know about the statistical aspects of genotype × environment interactions? Crop Science, 56(5), 2119–2140. 10.2135/cropsci2015.06.0375

Vergara, B. S., & Chang, T.-T. (1985). The flowering response of the rice plant to photoperiod: A review of the literature.

Via, S., Gomulkiewicz, R., Jong, G. D., Scheiner, S. M., Schlichting, C. D., & Tienderen, P. H. V. (1995). Adaptive phenotypic plasticity: Consensus and controversy. Trends in Ecology & Evolution, 10(5), 212–217. 10.1016/S0169-5347(00)89061-8

Went, F. (1953). The effect of temperature on plant growth. Annual Review of Plant Physiology, 4(1), 347–362.

Zhao, L., Li, M., Xu, C., Yang, X., Li, D., Zhao, X., Wang, K., Li, Y., Zhang, X., Liu, L., Ding, F., Du, H., Wang, C., Sun, J., & Li, W. (2018). Natural variation in GmGBP1 promoter affects photoperiod control of flowering time and maturity in soybean. The Plant Journal, 96(1), 147–162. 10.1111/tpj.14025.

